# Each cell needs long double-stranded RNA for silencing by feeding RNAi in *C. elegans*

**DOI:** 10.1101/066415

**Authors:** Pravrutha Raman, Soriayah M Zaghab, Edward C Traver, Antony M Jose

**Affiliations:** Department of Cell Biology and Molecular Genetics, University of Maryland, College Park, MD-20742; Present address: University of Maryland School of Medicine, Baltimore, MD 21201; Present address: Hofstra Northwell School of Medicine, Hofstra University, Hempstead, NY 11549

## Abstract

Long double-stranded RNA (dsRNA) can silence genes of matching sequence upon ingestion in many invertebrates and is therefore being developed as a pesticide. Such feeding RNA interference (RNAi) is best understood in the worm *C. elegans*, where it is thought that derivatives of ingested dsRNA, including short dsRNAs, move between cells and cause systemic silencing. Movement of short dsRNAs has been inferred using tissue-specific rescue of the long dsRNA-binding protein RDE-4 by expressing it from repetitive transgenes. We found that the use of repetitive transgenes for the tissue-specific rescue of a gene could inhibit RNAi within that tissue and could result in misexpression of the gene in other tissues. Both inhibition and misexpression were not detectable when a single-copy transgene was used for tissue-specific rescue. In animals with single-copy rescue of RDE-4, RNAi was restricted to the tissue with RDE-4 expression. Thus, unlike previous observations using repetitive transgenes, these results suggest that binding of long dsRNA by RDE-4 in each silenced cell is required for systemic RNAi. Taken together with the requirement for long dsRNA to trigger RNAi in insects, these results suggest that the entry of long dsRNA is a necessary first step for feeding RNAi in animal cells.

## INTRODUCTION

Killing animals by feeding them double-stranded RNA (dsRNA) that matches an essential gene is a powerful way to control animal pests. For example, expression of long dsRNA in potato plants was recently used to control populations of Colorado potato beetle that feed on these plants (1). This approach to pest control relies on the ability of many insects and parasitic nematodes to process ingested long dsRNA and use it to silence genes of matching sequence through RNA interference (RNAi) (2,3, reviewed in 4). However, the mechanisms of gene silencing by ingested dsRNA are not well understood, making it difficult to anticipate resistance mechanisms and therefore design effective dsRNA pesticides.

Silencing of genes by feeding animals long dsRNA was first demonstrated in the nematode *C. elegans* (5) and it remains the best animal model for understanding this process called feeding RNAi. In *C. elegans*, ingested dsRNA enters the animal through the intestine and can be delivered without entry into the cytosol of intestinal cells into the fluid-filled body cavity that surrounds all internal tissues (6-8). Entry into the cytosol of any cell requires a dsRNA-selective importer SID-1 (9) – a conserved protein with homologs in many insects (10). Upon entry into cells, silencing by dsRNA is thought to occur through the canonical RNAi pathway (reviewed in 11). Long dsRNA is first bound by the dsRNA-binding protein RDE-4, which recruits the endonuclease DCR-1 to generate short dsRNAs (12-14). One strand of this short dsRNA duplex is used as a guide by the primary Argonaute RDE-1 to identify mRNAs of matching sequence (15,12) and to recruit RNA-dependent RNA polymerases (RdRPs) to the mRNA. RdRPs then synthesize numerous secondary small RNAs (16-18) that are used for potent gene silencing within the cytosol by cytosolic Argonautes and/or within the nucleus by nuclear Argonautes (16-22). To infer whether any derivatives of long dsRNA generated within a cell also move between cells, components of the RNAi pathway were rescued in a specific tissue using repetitive transgenes and silencing was assayed in other tissues (23) or in the next generation (24). These tissue-specific rescue experiments suggested that short dsRNAs could be transported from donor cells to initiate gene silencing independent of RDE-4 in recipient cells.

Here, we demonstrate limitations of using repetitive transgenes which have resulted in erroneous inferences about feeding RNAi and that the use of single-copy transgenes can overcome these limitations. Unlike when repetitive transgenes were used, when single-copy transgenes were used for tissue-specific rescue of RDE-4, silencing was restricted to cells that expressed RDE-4. This requirement for RDE-4 suggests that all cells need the entry of long dsRNA for silencing in response to feeding RNAi.

## MATERIALS AND METHODS

### Worm Strains

All strains were cultured on Nematode Growth Medium (NGM) plates seeded with 100 μl OP50 at 20°C and mutant combinations were generated using standard methods (25). All strains used are listed in Supplemental Material.

### Balancing loci

Integrated transgenes expressing *gfp* were used to balance mutations in heterozygous animals. Progeny of heterozygous animals were scored as homozygous mutants if they lacked both copies of the transgene. The *rde-4(ne301)* allele on Chr III was balanced by *juls73*. About 99% (153/155) of the progeny of *rde-4(ne301)/juls73* that lacked fluorescence were found to be homozygous *rde-4(ne301)* animals either by Sanger sequencing (n = 96) or by resistance to *pos-1* RNAi (n = 59).

### Transgenesis

To make strains, N2 gDNA was used (unless otherwise specified) as a template to amplify promoter or gene regions. To amplify *gfp* to be used as a coinjection marker, a plasmid containing *gfp* sequence was used as a template (unless otherwise specified). All PCRs were performed with Phusion Polymerase (New England Biolabs - NEB), unless otherwise specified, according to the manufacturer’s recommendations. The final fusion products were purified using PCR Purification Kit (QIAquick, Qiagen).

### Plasmids

The plasmid pJM6 (made by Julia Marré, Jose lab) was used to make *Si[Pnas-9::rde-4(+)::rde-4 3’UTR]*. The *nas-9* promoter (*Pnas-9*) was amplified using primers P48 and P49 and *rde-4(+)::rde-4 3’UTR* was amplified using primers P50 and P4. The two PCR products were used as templates to generate the *Pnas-9::rde-4(+)::rde-4 3’UTR* fusion product with primers P51 and P52. This fused product was purified and cloned into pCFJ151 using the SbfI and SpeI restriction enzymes (NEB) to generate pJM6.

The plasmid pPR1 was used to make *Si[Pmyo-3::rde-4(+)::rde-4 3’UTR]*. The *myo-3* promoter (*Pmyo-3*) was amplified using primers P77 and P78 and *rde-4(+)::rde-4 3’UTR* was amplified using primers P79 and P4. The two PCR products were used as templates to generate the *Pmyo-3::rde-4(+)::rde-4 3’UTR* fusion product with primers P80 and P81. This fused product was purified and cloned into pCFJ151 using the SbfI restriction enzymes (NEB) to generate pPR1.

The plasmid pYC13 (made by Yun Choi, Jose lab) is a derivative of pUC::unc-119_sgRNA with a different sgRNA (gift from John Calarco, Addgene plasmid #46169).

All other plasmids were as described earlier (pHC448 (23), pPD95.75 (gift from Andrew Fire, Addgene plasmid #1494), pBH34.21 (26), pCFJ151 (27), pCFJ601 (27), pMA122 (27), pGH8 (27), pCFJ90 (27), pCFJ104 (27), pL4440 (5), pHC183 (6), and pGC306 (a gift from Jane Hubbard, Addgene plasmid #19658)).

### Genome editing

To generate *bli-1* null mutants, Cas9-based genome editing using a co-conversion strategy was used (28). Guide RNA for *bli-1* was amplified from pYC13 using primers P55 and P56, and guide RNA for co-conversion of *dpy-10* was amplified from pYC13 using P57 and P56. The amplified guides were purified (PCR Purification Kit, Qiagen) and tested *in vitro* for cutting efficiency (Cas9, NEB). For injection into animals, homology template for repair (repair template) was amplified from N2 gDNA using Phusion polymerase and gene specific primers. P58 and P59 were used to amplify a region immediately upstream of the 5’ region of *bli-1* and P60 and P61 were used to amplify a region immediately downstream of the 3’ region of *bli-1* using Phusion Polymerase (NEB). The two PCR products were used as templates to generate the repair template with primers P62 and P63 using Phusion Polymerase (NEB) and the fused product was purified (PCR Purification Kit, Qiagen). Homology template for *dpy-10* was a single-stranded DNA oligo (P64). Wild-type animals were injected with 3.5pmol/μl of *bli-1* guide RNA, 2.4pmol/μl of *dpy-10* guide RNA, 0.06pmol/μl of *bli-1* homology repair template, 0.6pmol/μl of *dpy-10* homology repair template and 1.6pmol/μl of Cas-9 protein (NEB). Resulting progeny animals were analyzed as described in Supplementary Figure S4.

### Feeding RNAi

RNAi experiments were performed on RNAi plates (NGM plates supplemented with 1 mM IPTG (Omega Bio-Tek) and 25μg/ml Carbenicillin (MP Biochemicals)) at 20°C.

#### One generation or F1-only feeding RNAi

A single L4 or young adult (1 day older than L4) animal (P0) was placed on an RNAi plate seeded with 5μl of OP50 *E. coli* and allowed to lay eggs. After 1 day, by when most of the OP50 *E. coli* was eaten, the P0 animal was removed, leaving the F1 progeny. 100μl of an overnight culture of RNAi food (*E. coli* which express dsRNA against a target gene) was added to the plate. Two or three days later, the F1 animals were scored for gene silencing by measuring gene-specific defects (See Supplementary Table S1). All RNAi *E. coli* clones were from the Ahringer library (29) and generously supplied by Iqbal Hamza, with the exception of *unc-54* RNAi, which was made by cloning a fragment of *unc-54* DNA into pL4440 and transforming into HT115(DE3) *E. coli* cells. Control RNAi by feeding *E. coli* containing the empty dsRNA-expression vector (pL4440), which does not produce dsRNA against any gene, was done in parallel with all RNAi assays.

Fluorescence intensity of L4 animals fed dsRNA against *gfp* was measured using Image J (National Institute of Health-NIH). Animals with an intensity of >6000 (a.u.) in a fixed area within the gut immediately posterior to the pharynx were considered not silenced. Based on this criteria, 91.7% of wild-type animals and a 100% of animals expressing *DsRed* in the muscle (*Ex[Pmyo-3::DsRed]*) showed silencing. In these silenced animals intensity of *gfp* was measured from below the pharynx to the end of the vulva and the background intensity for the same area was measured for each animal. The intensity of *gfp* for the area after background subtraction was plotted for each worm (Supplementary Figure S2D).

#### Two generations or P0 & F1 Feeding RNAi

The experiments in all Figures (except Figure 2, Supplementary Figures S1B and C, S5A and B, and S6) were performed by feeding both the P0 and F1 generations, as described earlier (6). Control RNAi was done in parallel with all RNAi assays. Three or four days after P0 animals were subjected to RNAi, the F1 animals were scored for gene silencing by measuring gene-specific defects (See Supplementary Table S1).

**Figure 1.**
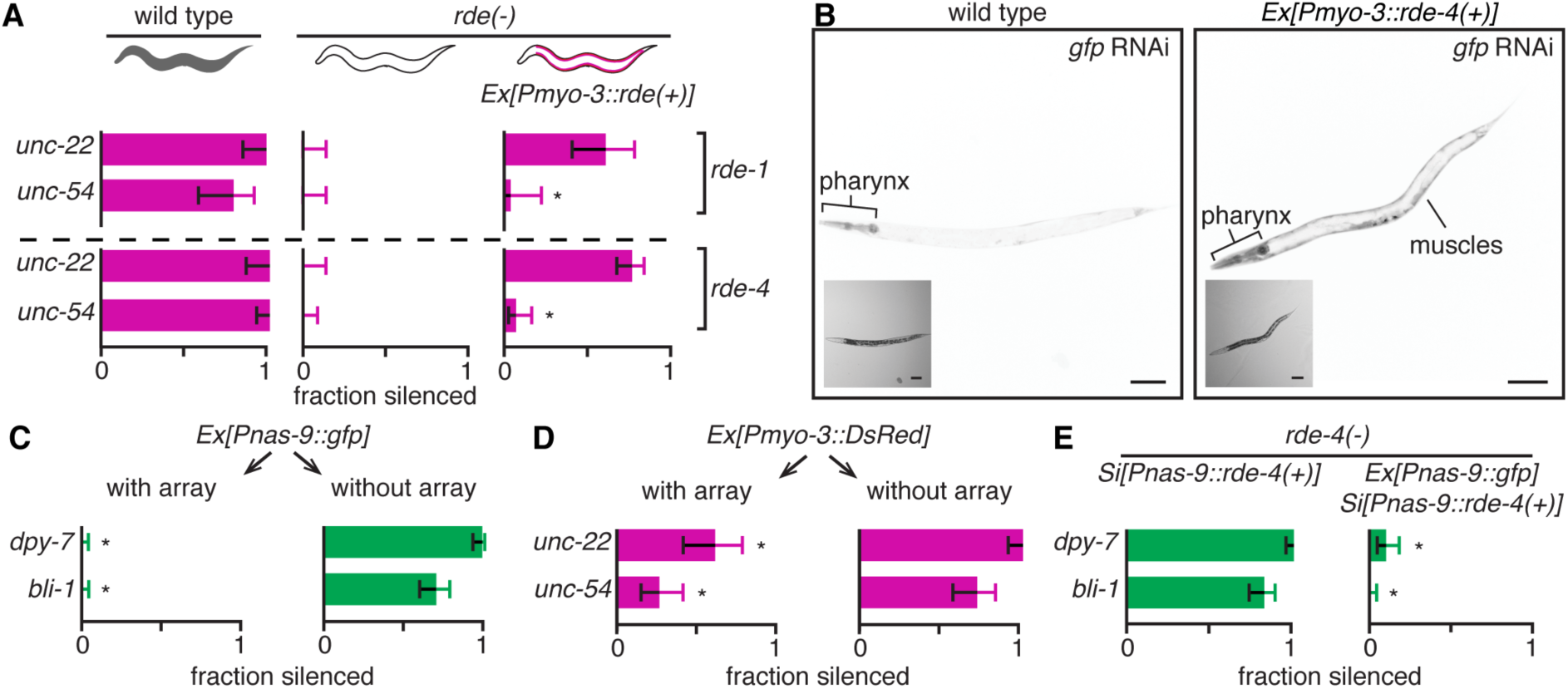
Expression of any repetitive transgene in a tissue can inhibit silencing by ingested dsRNA within that tissue. (**A**) Silencing by feeding RNAi of some endogenous genes is reduced in tissues expressing *rde-4(+)* or *rde-1(+)* from a repetitive transgene. Wild-type animals, mutant animals (*rde-1(-)*, top or *rde-4(-)*, bottom) or mutant animals with tissue-specific rescues in the body-wall muscles (*Ex[Pmyo-3::rde(+)]*) were fed dsRNA against *unc-22* or *unc-54* and the fractions of animals that showed silencing (fraction silenced) were determined. (**B**) Silencing of *gfp* in body-wall muscles that express RDE-4 from a repetitive transgene is reduced despite potent silencing in other *rde-4(-)* soma. Representative images of animals with *gfp* expression in all somatic cells (*Peft-3::gfp*) in a wild-type background (left) or *rde-4(-)* background with *rde-4(+)* expressed in body-wall muscles (*Ex[Pmyo-3::rde-4(+)*) (right) that were fed bacteria that express dsRNA against *gfp* (*gfp* RNAi) are shown. Tissues that show reduced silencing (pharynx, muscles) are labelled, insets are brightfield images, and scale bar = 50 μm. Also see Supplementary Figure S1B and C. (**C**and **D**) Silencing of a gene by ingested dsRNA within a tissue can be inhibited by the expression of a repetitive transgene of unrelated sequence within that tissue. (C) Silencing of *bli-1* and *dpy-7* is inhibited by the expression of *gfp* from a repetitive transgene in the hypodermis. Wild-type animals that express *gfp* alone in the hypodermis (*Ex[Pnas-9::gfp]*) from extrachromosomal repetitive DNA (array) were fed dsRNA against hypodermal genes (*dpy-7* or *bli-1*, green). The fractions of animals either with or without the arrays that showed silencing (fraction silenced) were determined. (D) Silencing of *unc-22* and *unc-54* can be inhibited by the expression of *DsRed* from a repetitive transgene in body-wall muscles. Wild-type animals expressing *DsRed* in the body-wall muscle (*Ex[Pmyo-3::DsRed]*) from extrachromosomal repetitive DNA (array) were fed dsRNA against body-wall muscle genes (*unc-22* or *unc-54*, magenta). Silencing was determined as in (C). (**E**) Expression of RDE-4 from a single-copy transgene within a tissue does not inhibit feeding RNAi in that tissue. *rde-4(-)* animals that express RDE-4 in the hypodermis from a single-copy transgene (*Si[Pnas-9::rde-4(+)]*) or that additionally express *gfp* in the hypodermis (*Ex[Pnas-9::gfp]*) from a repetitive transgene (array) were fed dsRNA against *dpy-7* or *bli-1* and were analyzed as in (C). Error bars indicate 95% confidence intervals (CI), n>23 animals, and asterisks indicate p<0.01. Also see Supplementary Figures S1 and S2.

**Figure 2.**
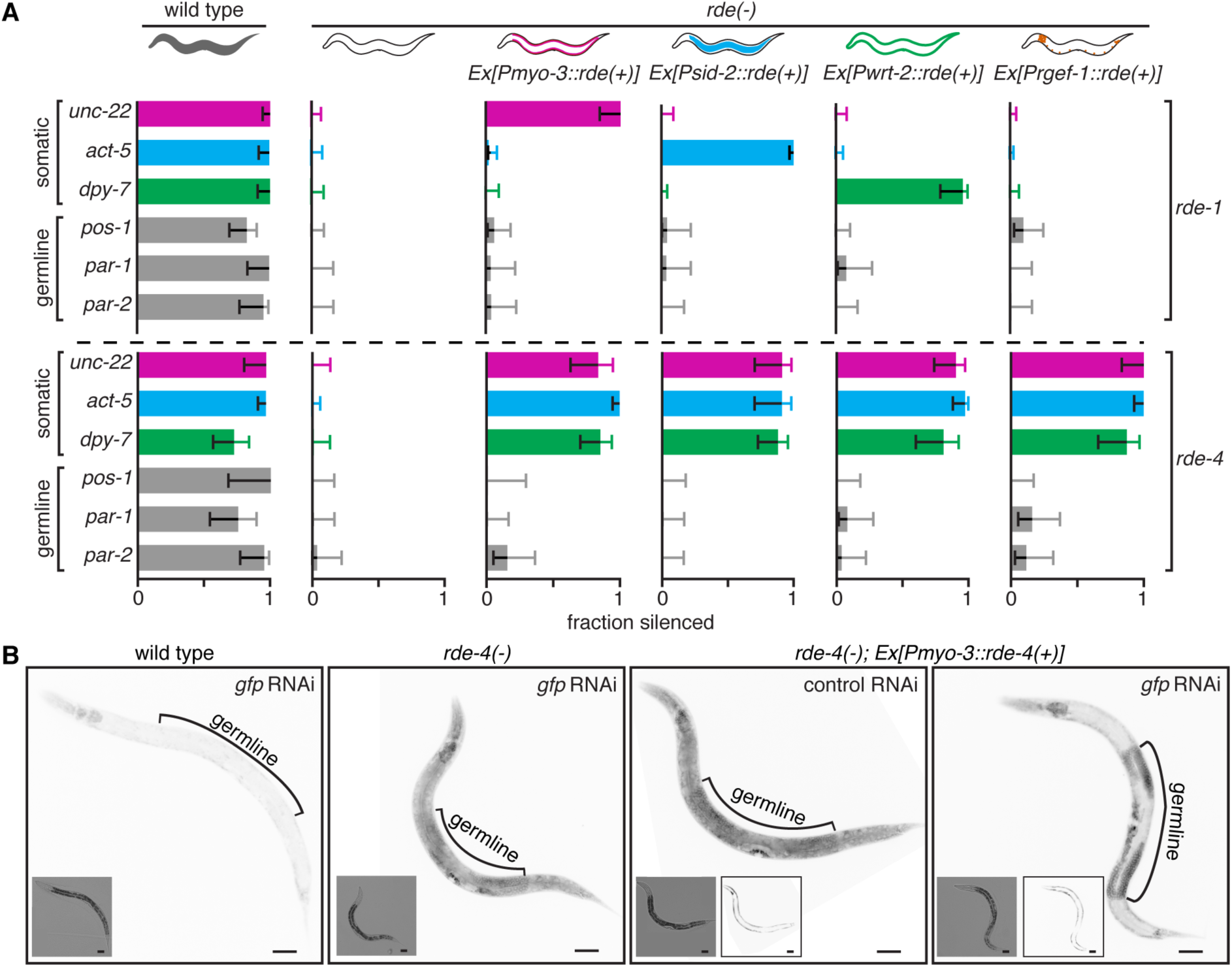
Tissue-specific rescues of RDE-4 from repetitive transgenes can enable silencing of genes in mutant somatic tissues. (**A**) Tissue-specific expression of RDE-4 but not RDE-1 from repetitive transgenes enables silencing of endogenous genes that function in mutant somatic tissues but not in mutant germline. Wild-type animals (grey), mutant animals (*rde-1(-)* or *rde-4(-)*, white), and mutant animals with tissue-specific rescues (colors within worms) of *rde-1* or *rde-4* were fed dsRNA against genes expressed in somatic tissues (the body-wall muscles (*unc-22*, magenta), intestine (*act-5*, blue), hypodermis (*dpy-7*, green)), or in the germline (*pos-1, par-1*, or *par-2*, grey) and the fractions of animals that showed silencing (fraction silenced) were determined. Wild-type genes (*rde-1* or *rde-4*) were expressed in the body-wall muscles (*Ex[Pmyo-3::rde(+)]*, magenta), in the intestine (*Ex[Psid-2::rde(+)]*, blue), in the hypodermis (*Ex[Pwrt-2::rde(+)]*, green), or in neurons (*Ex[Prgef-1::rde(+)]*, orange). Error bars indicate 95% CI and n>24 animals. (**B**) Tissue-specific rescue of *rde-4* enables silencing of *gfp* in *rde-4(-)* somatic tissue but not in *rde-4(-)* germline. Representative images of animals with *gfp* expression in all somatic and germline cells (*Pgtbp-1::gtbp-1::gfp*) in a wild-type background, *rde-4(-)* background, or *rde-4(-)* background with *rde-4(+)* expressed in body-wall muscles (*Ex[Pmyo-3::rde-4(+)*) that were fed control RNAi or *gfp* RNAi are shown. In all cases, 50 L4-staged animals were analysed and the majority phenotype (wild-type – 100%, *rde-4(-)* – 100%, *rde-4(-);Ex[Pmyo-3::rde-4(+)* control RNAi *-* 100%, and *rde-4(-);Ex[Pmyo-3::rde-4(+) gfp* RNAi – 68%) is shown. Insets are brightfield images and scale bar = 50 μm.

No difference in gene silencing was observed between F1-only feeding RNAi and P0 & F1 feeding RNAi for *rde-4* mutants with tissue-specific rescue. For each RNAi experiment testing *rde-4* function, feeding of N2 and WM49 was performed alongside as controls. In the case of Supplementary Figure S7 (*act-5* feeding RNAi of *rde-4(-); Si[nas-9::rde-4(+)::rde-4 3’UTR; Ex[Pnas-9::gfp::unc-54 3’UTR]*), L4 animals were scored a day after control L4 animals because these animals grew slower than control animals.

### Scoring defects

For RNAi treatments, the proportions of animals that displayed the reported mutant defects upon RNAi (see Supplementary Table S1) were scored as “fraction silenced”.

For *bli-1* defects upon RNAi and upon Cas9-based genome editing, the pattern of blister formation was scored. Each animal was partitioned into eight roughly equal sections (a to h) as shown in Figure 3C with the vulva being the mid-point of the animal. Sections with >50% of their length covered by a blister were marked black and sections with <50% of their length covered in a blister or with a discontinuous blister were marked grey. Animals that did not follow the anterior more than posterior and dorsal more than ventral susceptibility pattern (a > b > c > d > e > f > g > h) were culled as variants for each genotype and the relative aggregate blister formation in each section among worms with altered susceptibility (Figure 3D, Supplementary Figure S4E-H) were computed using a score of black = 1.0 and grey = 0.5 for each section of every worm. The computed values for each section in all worms of a strain were summed and normalized to the value of the highest section for that strain. To compare multiple strains, these values for each strain were multiplied by the fraction of worms that showed a blister in that strain. Using these measures of normalized relative aggregate blister formation among animals with variant susceptibility, we generated heat maps (30), where black indicates highest frequency of blisters and white indicates the lowest frequency of blisters among the sections of all strains that are being compared.

**Figure 3.**
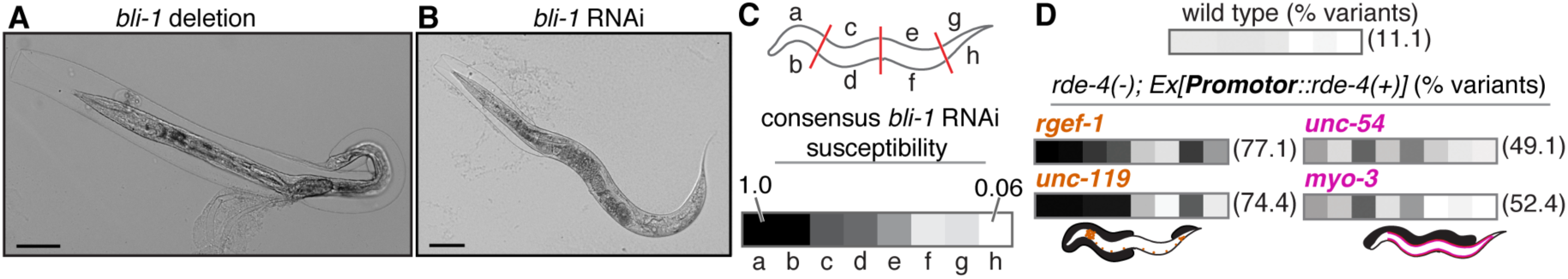
Repetitive transgenes expressing RDE-4 from different promoters enable different spatial patterns of *bli-1* silencing in *rde-4(-)* hypodermis. (**A**) A null mutation in *bli-1* results in blisters that cover the entire body. A representative image of a *bli-1* null mutant animal generated by Cas9-based genome editing. Scale bar = 50 μm. (**B**) Feeding RNAi of *bli-1* in wild-type animals results in blisters that cover part of the body. A representative image of a wild-type animal fed dsRNA against *bli-1* (*bli-1* RNAi). Scale bar = 50 μm. (**C**) Susceptibility to *bli-1* feeding RNAi decreases from anterior to posterior hypodermis in wild-type animals. (top) Schematic of hypodermal sections (a through h) scored for blister formation. (bottom) Consensus relative frequency of blister formation in each hypodermal section of wild-type animals upon *bli-1* feeding RNAi. The frequency ranged from 1.0 (black, a) to 0.06 (~white, h). (**D**) The patterns of blisters that result from silencing of *bli-1* in *rde-4(-)* hypodermis are different from the consensus blister pattern. Aggregate patterns of blister formation among animals that deviate from the consensus susceptibility order (consensus *bli-1* RNAi susceptibility in (C)) for each strain (variant susceptibility, % variants) are shown. All strains being compared were normalized together (black, section with highest frequency of blisters in all strains; white, section with the lowest frequency of blisters in all strains). Schematic of worms indicate locations of variant blisters (thick black shading) on worms with *rde-4(+)* expressed in neurons (orange) or in body-wall muscles (magenta). Also see Supplementary Figure S4.

### Microscopy

Animals were immobilized in 5μl of 3mM levamisole, mounted on slides, and imaged using an AZ100 microscope (Nikon) at a fixed magnification under non-saturating conditions. Images being compared on any figure were adjusted identically using Adobe Photoshop (levels adjustment) for display.

ImageJ (NIH) was used to generate merged images (Supplementary Figure S2A-C). To generate merged images, the LUT was set from 0 (white) to 127 (magenta) for DsRed and from 0 (white) to 127 (green) for GFP. One channel was then overlayed on the other with 50% opacity.

### Statistical Analyses

Error bars in all cases indicate 95% confidence intervals for single proportions calculated using Wilson’s estimates with a continuity correction (Method 4 in (31)). Significance of differences between two strains or conditions was determined using pooled Wilson’s estimates.

### Data Availability

All strains are available upon request.

## RESULTS

### Expression of a repetitive transgene in a tissue can inhibit RNAi in that tissue

Until recently, studies examining the function of *C. elegans* genes have relied on the use of repetitive transgenes often coupled with tissue-specific promoters (e.g. 32-34) and/or RNAi (e.g. 35-37). Such an approach was also used to examine silencing by feeding RNAi. Rescue of *rde-4* and *rde-1* using the *myo-3* promoter to drive expression in body-wall muscles from a repetitive transgene (*Ex[Pmyo-3::rde(+)]*) was used to examine the systemic response to RNAi (23,24). While silencing in *rde-1(-)* animals with *Ex[Pmyo-3::rde-1(+)]* was observed only in body-wall muscles, silencing in *rde-4(-)* animals with *Ex[Pmyo-3::rde-4(+)]* was observed in both body-wall muscles and other tissues. We found that in both cases silencing of some genes within body-wall muscle cells was substantially reduced compared to that in wild-type animals in response to feeding RNAi (compare *unc-22* silencing versus *unc-54* silencing in Figure 1A). Similar reduction in silencing was observed in hypodermal cells upon hypodermal rescue of *rde* genes (compare *dpy-7* silencing versus *bli-1* silencing in Supplementary Figure S1A). When a ubiquitously expressed target gene was tested in *rde-4(-); Ex[Pmyo-3::rde-4(+)]* animals, this reduction of silencing was observed only within body-wall muscles despite robust silencing in other tissues (*gfp* silencing in Figure 1B and Supplementary Figure S1B and C). Extent of silencing of a gene varied based on the specific promoter used (e.g. silencing of *dpy-7* when *wrt-2* promoter was used (Supplementary Figure S1A) versus when *nas-9* promoter was used (Supplementary Figure S1D)). Thus, rescuing *rde-4* or *rde-1* using tissue-specific promoters and repetitive transgenes does not reliably restore tissue-restricted silencing of all genes.

To test if these cases of reduced silencing could be explained by insufficient levels of *rde* expression, we overexpressed RDE-4 in the hypodermis of wild-type animals (*Ex[Pnas-9::rde-4(+)]*) and examined silencing. Feeding RNAi of neither *bli-1* nor *dpy-7* resulted in detectable silencing (Supplementary Figure S1D). Thus, the observed lack of silencing is not because there was insufficient *rde-4(+)* expression but because expression of *rde-4(+)* within a tissue from a repetitive transgene inhibited feeding RNAi in that tissue. Such lack of silencing for both tested genes could reflect co-suppression (38) of *rde-4*. However, this hypothesis cannot explain the differential susceptibility of *bli-1* and *dpy-7* that was observed when *rde-4(+)* was expressed under the *wrt-2* promoter. Alternatively, expression of any repetitive transgene could inhibit RNAi and the extent of inhibition could vary based on the target gene being tested and the promoter used. To test these possibilities, we expressed *gfp* from a repetitive transgene in the hypodermis (*Ex[Pnas-9::gfp]*) and examined silencing of *bli-1* and of *dpy-7* by feeding RNAi. Surprisingly, no silencing was detected when *gfp* was expressed (Figure 1C), suggesting that inhibition of silencing is the result of expression from a repetitive transgene and not because of *rde-4* expression or co-suppression. Similar inhibition of silencing of *unc-54, unc-22*, and *gfp* was also observed in body-wall muscles when we expressed *DsRed* from repetitive transgenes in body-wall muscles (Figure 1D, Supplementary Figure S2). Together, our results suggest that expression from repetitive transgenes in a tissue can interfere with silencing by ingested dsRNA within that tissue.

Repetitive transgenes can produce dsRNA (39,40) that could compete with ingested dsRNA for engaging the gene silencing machinery within a cell. Such competition between pathways is the reason silencing by feeding RNAi can be enhanced in animals that lack genes required solely for the processing of endogenous dsRNA (41,42, reviewed in 43). Consistently, we found that loss of factors required for endogenous dsRNA production (the RdRP *rrf-3* or the endonuclease *eri-1*) but not loss of downstream Argonaute proteins (the primary Argonaute *ergo-1* or the secondary Argonaute *nrde-3*) overcame the inhibition of feeding RNAi (Supplementary Figure S3A, right). Because RRF-3 and ERI-1 are not required for the production of dsRNA from repetitive transgenes (44), these results suggest that silencing by feeding RNAi in the presence of expression from a repetitive transgene is enabled by loss of dsRNA production at endogenous loci (see Supplementary Figure S9 in 45).

Certain genes appear to be more susceptible to inhibition by expression from repetitive transgenes. For example, *unc-54* was more susceptible than *unc-22* (Figure 1A) and *bli-1* was more susceptible than *dpy-7* (Supplementary Figure S1A). While the basis for these differences is unclear, we found silencing of *bli-1* and *unc-54* but not of *unc-22* showed a dependence on the secondary Argonaute NRDE-3 (Supplementary Figure S3B). Consistent with a role for the nuclear RNAi pathway in silencing these genes, we found that *bli-1* silencing by feeding RNAi also depended on components that act downstream of NRDE-3 (46,21) (Supplementary Figure S3C). This dependence on the nuclear RNAi pathway was also observed when *bli-1* was targeted for silencing by dsRNA expressed in neurons (Supplementary Figure S3D), suggesting that irrespective of the source of the dsRNA, NRDE-3 is required for silencing *bli-1*. Additional experiments are needed to establish mechanistic links, if any, between *nrde-3*-dependence and inhibition of RNAi by expression from repetitive transgenes.

Taken together, our analysis predicts that the use of a single-copy transgene should eliminate the inhibition observed and enable silencing by RNAi. Accordingly, using a single-copy transgene to express *rde-4(+)* in the hypodermis (*Si[Pnas-9::rde-4(+)]*) enabled potent silencing of both *dpy-7* and *bli-1* by feeding RNAi (Figure 1E, left). Furthermore, this silencing could be inhibited by additionally expressing *Ex[Pnas-9::gfp]* (Figure 1E, right). Thus, expression of any repetitive transgene in a tissue can inhibit silencing of some genes within that tissue and using a single-copy transgene can avoid this problem.

### Rescue of *rde-4* in one somatic tissue from repetitive transgenes can cause silencing within most somatic tissues but not in the germline

When repetitive transgenes were used to rescue *rde-4* in body-wall muscles, silencing upon feeding RNAi was observed in other somatic tissues (23). In contrast, when repetitive transgenes were used to rescue *rde-1* in body-wall muscles, silencing was restricted to the tissue where *rde-1* was expressed (47,23). These results were used to infer the intercellular movement of possibly short dsRNAs generated downstream of RDE-4 but upstream of RDE-1. To test if this observation extended to other tissues (intestine, hypodermis, and neurons), we similarly used repetitive transgenes to perform tissue-specific rescues of *rde-1* and *rde-4* and assessed silencing of genes expressed in somatic tissues and the germline (Figure 2A). In all cases, silencing by feeding RNAi was observed only in one RNAi-sensitive somatic tissue when RDE-1 was expressed in that somatic tissue (Figure 2A, top) but in all tested somatic tissues when RDE-4 was expressed in any one somatic tissue (Figure 2A, bottom). In contrast, no silencing was detectable within the germline in any case (Figure 2A). This difference between silencing in soma and germline was also observed when the same ubiquitously expressed gene (*Pgtbp-1::gtbp-1::gfp*) was targeted for silencing in *rde-4(-); Ex[Pmyo-3::rde-4(+)]* animals (Figure 2B). Together, these results could be interpreted as supporting either misexpression of RDE-4 in unintended somatic tissues in all cases or transport of short dsRNAs between somatic cells when long dsRNA is processed in one somatic tissue by RDE-4.

### Spatial patterns of silencing vary with the promoters that drive tissue-specific rescue from repetitive transgenes

If transport of short dsRNAs, rather than misexpression, is the reason for the observed silencing, then silencing could be more common in cells that are near the source of dsRNA. However, when an animal is only scored as silenced versus not silenced in response to feeding RNAi, such qualitative differences between animals are overlooked. Examination of such differences requires a target gene whose silencing in subsets of cells can be discerned in each animal. We found that null mutants of the hypodermal gene *bli-1* result in a fluid-filled sac (“blister”) along the entire worm (Figure 3A, Supplementary Figure S4A and B; (25)), and that blisters that form upon feeding RNAi in wild-type animals had a different pattern (Figure 3B). Specifically, anterior sections of the worm tended to be more susceptible to silencing when compared with posterior sections (Figure 3C, Supplementary Figure S4C, and see methods), resulting in a stereotyped pattern of relative susceptibility to blister formation upon *bli-1* feeding RNAi (Figure 3C, bottom and Supplementary Figure S4D). This bias in the tendency to form blisters likely reflects the graded uptake of dsRNA from the anterior to the posterior in the intestine upon feeding RNAi. These characteristics of blister formation as a result of *bli-1* silencing enable examination of qualitative differences, if any, between silencing in wild-type animals and in animals with tissue-specific *rde-4* rescue.

To systematically analyze such differences, we culled animals that had a pattern of blister formation that differed from a consensus blister susceptibility pattern observed in most wild-type animals (see methods). We found that unlike in wild-type animals, in animals with tissue-specific rescue of *rde-4* from repetitive transgenes, patterns of blisters that differed from the reference pattern were common (Figure 3D and Supplementary Figure S4E-H). Furthermore, the pattern of variant blister susceptibility differed depending on the promoter used for tissue-specific rescue (*rgef-1* or *unc-119* for neurons and *myo-3* or *unc-54* for body-wall muscles) (Figure 3D and Supplementary Figure S4G and H). Intriguingly, animals with different promoters that drive expression in the same tissue showed similar patterns of silencing. While these results could provide a case for the transport of short dsRNAs to nearby cells from the tissue where long dsRNA is processed by RDE-4, similar results would also be obtained if misexpression from each tissue-specific promoter occurred in subsets of hypodermal cells.

### Silencing within a somatic tissue requires RDE-4 in that tissue

Consideration of the following observations on repetitive transgenes further support the possibility that misexpression of *rde-4(+)* in somatic tissues explains the silencing in all somatic tissues despite the use of tissue-specific promoters. First, the formation of a repetitive transgene can generate rearrangements that result in novel promoter elements (48,49,40) that could lead to misexpression despite the use of well-characterized tissue-specific promoters. In support of this possibility, in animals expressing *gfp* in all somatic tissues (*Peft-3::gfp*) some silencing in non-muscle cells was detected even in *rde-1(-); Ex[Pmyo-3::rde-1(+)]* animals (Supplementary Figure S5). Second, repetitive transgenes can be selectively silenced within the germline (50,51), potentially explaining the observed lack of silencing in the germline of animals with tissue-specific *rde-4* rescue (Figure 2). Third, because repetitive transgenes could be expressed at low levels within the germline (e.g. heat shock promoter (52)), inherited RDE-4 from such expression could support feeding RNAi in all somatic tissues as was observed in *rde-4(-)* progeny of heterozygous parents (53). While such silencing enabled by inherited RDE-4 was not detectable in most cases, it was detectable when *unc-22* silencing was examined in *rde-4(-)* progeny of animals with *rde-4* rescued using the *myo-3* promoter (Supplementary Figure S6).

We attempted to distinguish between the movement of dsRNA between cells and the presence of RDE-4 in unintended somatic tissues upon tissue-specific rescue by restricting the presence of the dsRNA-selective importer SID-1 (9,54,55). Tissues that express high levels of SID-1 act as sinks for dsRNA (7) and the entry of both long dsRNA and short dsRNAs generated upon processing by RDE-4 are expected to require SID-1. We generated *sid-1(-); rde-4(-)* animals in which *rde-4* was rescued using the neuronal promoter *rgef-1* and *sid-1* was rescued using the body-wall muscle promoter *myo-3* (Figure 4A). In these animals, the entry of ingested dsRNA is expected to occur only into body-wall muscle cells (6-8). If expression from the *rgef-1* promoter only resulted in the presence of RDE-4 in neurons, then no silencing would be detected because while long dsRNA can enter body-wall muscles through SID-1, it cannot be processed further for silencing in the absence of RDE-4. On the other hand, if expression from the *rgef-1* promoter resulted in the presence of RDE-4 in neurons and in body-wall muscles, then silencing would be observed because dsRNA can both enter into and be processed in body-wall muscles. We observed silencing in these animals upon feeding RNAi of the body-wall muscle gene *unc-54* (Figure 4B). Consistent with the restriction of SID-1-dependent dsRNA entry into body-wall muscle cells, we did not detect any silencing of the hypodermal gene *bli-1*. Together these results suggest that misexpression of *rde-4(+)* in somatic tissues when repetitive transgenes are used or the movement of RDE-4 - but not dsRNA - between cells is sufficient to explain the silencing in all somatic tissues. Therefore, we re-examined if the expression of RDE-4 in one tissue could enable silencing in other somatic tissues using single-copy transgenes. We expressed RDE-4 from a single-copy transgene using the *myo-3* promoter (*Si[Pmyo-3::rde-4(+)]*) or the *nas-9* promoter (*Si[Pnas-9::rde-4(+)]*) in *rde-4(-)* animals. In both cases, silencing was restricted to the intended tissue with RDE-4 expression (Figure 4C). These results do not support the possibility that RDE-4 protein or mRNA moves between cells and suggest that short dsRNAs made in one tissue are not sufficient to cause silencing in another tissue.

**Figure 4.**
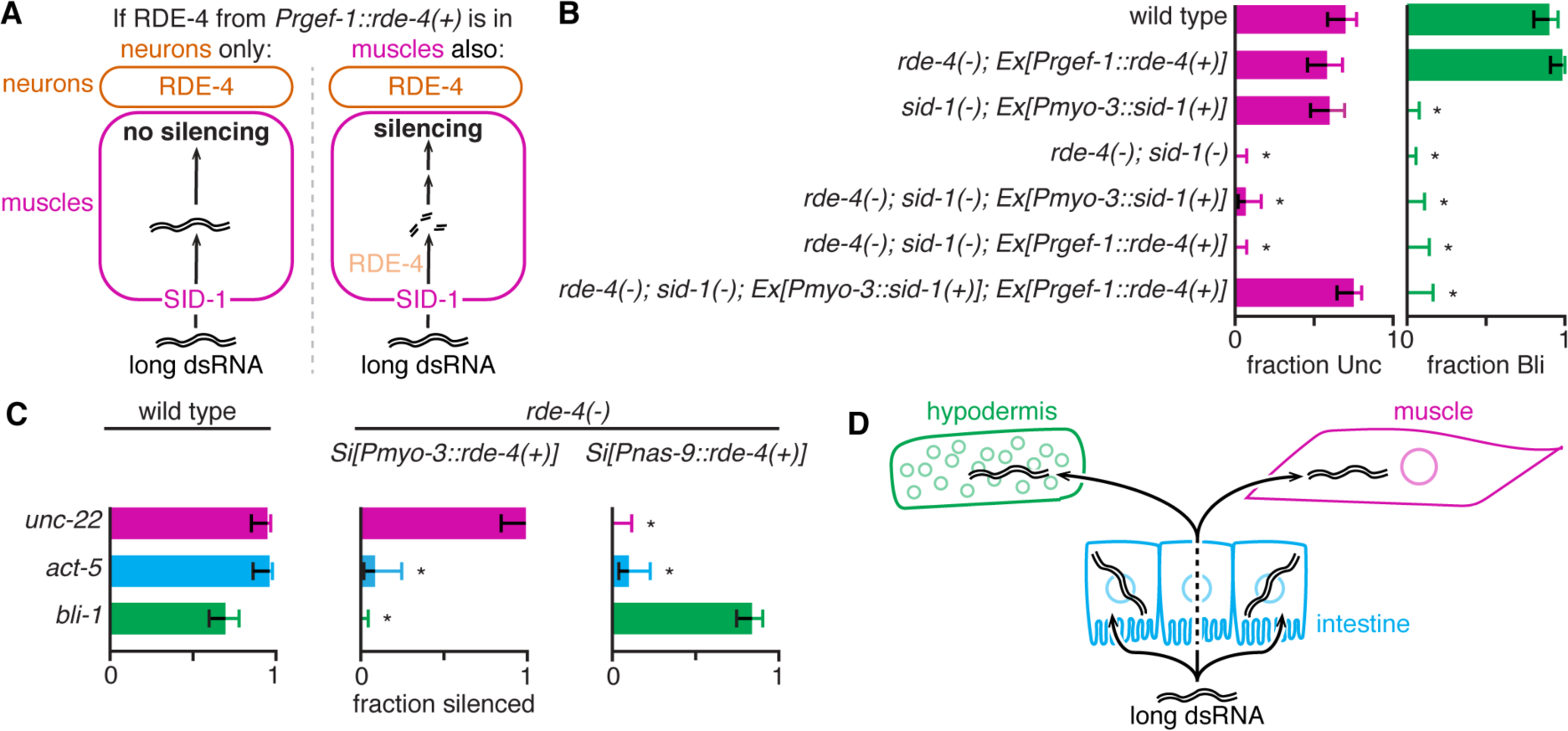
Silencing by ingested dsRNA in a tissue requires long dsRNA within that tissue. (**A**) Expected outcomes of test to distinguish movement of short dsRNAs from misexpression of RDE-4 in animals with tissue-specific rescue of RDE-4 from a repetitive transgene. (Left) If RDE-4 is expressed only in neurons from the *rgef-1* promoter (*Prgef-1::rde-4(+)*) in a *rde-4(-); sid-1(-); Ex[Pmyo-3::sid-1(+)]* background no silencing is expected in muscles, which can import dsRNA (have SID-1) but not process dsRNA (lack RDE-4). (Right) If RDE-4 is expressed in neurons and in muscles from the *rgef-1* promoter (*Prgef-1::rde-4(+)*) in a *rde-4(-); sid-1(-); Ex[Pmyo-3::sid-1(+)]* background silencing can occur in muscles, which can import dsRNA (have SID-1) and process dsRNA (have RDE-4). (**B**) RDE-4 is present in muscles when expressed under the *rgef-1* promoter from a repetitive transgene. Wild-type animals, mutant animals (*rde-4(-), sid-1(-)*, or *rde-4(-);sid-1(-)*) or mutant animals with *rde-4(+)* and/or *sid-1(+)* expressed from repetitive transgenes (as schematized in (A)) were fed dsRNA against a gene expressed in the body-wall muscle (*unc-54*) or in the hypodermis (*bli-1*) and the fractions of animals that showed silencing (fraction Unc or fraction Bli) were determined. No silencing was observed in *rde-4(-)* or in *sid-1(-)* animals. Error bars indicate 95% CI, n>24 animals and asterisks indicate p<0.01 (compared to wild-type animals). (**C**) Expression of RDE-4 from a single-copy transgene reveals a requirement for RDE-4 within a tissue for silencing by ingested dsRNA in that tissue. Wild-type animals or *rde-4(-)* animals that express RDE-4 from a single-copy transgene in the body-wall muscle (*Si[Pmyo-3::rde-4(+)]*) or in the hypodermis (*Si[Pnas-9::rde-4(+)]*) were fed dsRNA against *unc-22* (magenta), *act-5* (blue) or *bli-1* (green). Silencing was scored as in Figure 2A. Error bars indicate 95% CI, n>24 animals and asterisks indicate p<0.01. Also see Supplementary Figure S7. (**D**) Model: Ingested long dsRNA needs to enter each tissue to cause silencing in that tissue.

## DISCUSSION

Our analyses in *C. elegans* reveal that, contrary to previous models, systemic silencing by feeding RNAi requires entry of long dsRNA into all cells that show silencing (Figure 4D). We provide strong evidence that the previous erroneous models resulted from the use of repetitive transgenes to study gene function. Furthermore, repetitive transgenes can inhibit feeding RNAi selectively within the tissue where the transgene is expressed.

### Efficiency of RNAi could be regulated by expression from repetitive DNA

Our discovery that expression from repetitive DNA within a tissue can interfere with silencing by feeding RNAi within that tissue (Figure 1) could impact studies that use RNAi to infer the function of a gene. For example, RNAi of a gene in strains that express fluorescent reporters within a tissue from a repetitive transgene could be specifically inhibited in that tissue. RNAi screens performed on strains expressing repetitive transgenes could have missed genes that are sensitive to inhibition. Thus, inferences from feeding RNAi in strains with repetitive transgenes may need to be re-examined using single-copy transgenes.

The efficiency of feeding RNAi differs between tissues and is a key concern for the application of feeding RNAi to combat animal pests (4). For example, in *C. elegans*, genes expressed in neurons are relatively refractory to silencing by feeding RNAi (noted in (56)). One reason for such reduced silencing could be that neurons have high levels of expression from endogenous repetitive DNA. Consistent with this possibility, both silencing in tissues with expression from repetitive DNA (Supplementary Figure S3A) and silencing in neurons are enhanced upon loss of the exonuclease ERI-1 (57) or the RdRP RRF-3 (58). Similarly, tissue-specific expression from endogenous repetitive DNA could explain differential sensitivity to RNAi among insect tissues.

### Long dsRNA is needed in every cell for silencing when an animal is subjected to feeding RNAi

The ability of dsRNA expressed in one cell to cause SID-1-dependent silencing in other cells (9,6) revealed that dsRNA or its derivatives can be exported from cells and be imported into cells in *C. elegans*. The previous inference that derivatives of ingested dsRNA can also move between cells (23,24) resulted in a “transit” model for feeding RNAi where dsRNA first enters the cytosol of a cell, is subsequently processed within the cytosol of that cell, and finally exported for silencing in distant cells. Our discovery that RDE-4 is required in each cell for silencing upon feeding RNAi (Figure 4) does not support such a transit model, and yet accounts for all silencing upon feeding RNAi. Because dsRNA can be transported across intestinal cells without entry into the cytosol (6-8) and reach the pseudocoelom that bathes all *C. elegans* tissues, the direct entry of dsRNA into all cells that show silencing and subsequent processing by RDE-4 in each cell is sufficient to explain the systemic response to feeding RNAi.

Taken together with recent studies, our results suggest that several characteristics of feeding RNAi in many insects and parasitic nematodes (see 59,4,60 for reviews) could be similar to those in *C. elegans*. First, long dsRNA (>60 bp) is preferentially ingested (61) and realization of this preference was crucial for developing plastid expression as an effective strategy to deliver long dsRNA into crop pests (1). Second, dsRNA can be detected in intestinal cells and in internal tissues upon feeding RNAi (62). Third, with the exception of dipteran insects, most invertebrates have homologs of the dsRNA importer SID-1 (10). Finally, silencing initiated by feeding RNAi can persist for multiple generations (63). These similarities suggest that insights gleaned using the tractable animal model *C. elegans* are likely to be applicable to many invertebrates, including agronomically important insect and nematode pests.

## ACKNOWLEDGEMENT

We thank Leslie Pick and members of the Jose lab for critical reading of the manuscript; Julia Marre (Jose lab) for generating the plasmid pJM6; Yun Choi (Jose lab) for generating the plasmid pYC13; the *Caenorhabditis elegans* Genetic stock Center, the Hunter lab (Harvard University), and the Seydoux lab (Johns Hopkins University) for some worm strains and the Hamza lab (University of Maryland) for bacteria that express *gfp*-dsRNA. Critical comments from anonymous reviewers were crucial in arriving at the working model proposed in this manuscript.

## SUPPLEMENTAL MATERIALS AND METHODS

### Strains Used

N2 wild type

AMJ8 *juls73[Punc-25::gfp] III*

AMJ57 *sid-1(qt9) V; jamEx12[Pmyo-3::sid-1(+)::unc-54 3’ UTR & pHC183]* (strain generated in (6) after multiple passages)

AMJ58 *rde-1(ne219) I; jamEx1[Pmyo-3::rde-1(+)::rde-1 3’UTR & Pmyo-3::DsRed2::unc-54 3’UTR]*

AMJ66 *rde-4(ne301); jamEx4[Pmyo-3::rde-4(+)::rde-4 3’UTR & pHC183]* (strain generated in (23) after multiple passages).

AMJ151 *rde-4(ne301); mIs11[Pmyo-2::gfp::unc-54 3’UTR & gut::gfp::unc-54 3’UTR & pes-10::gfp::unc-54 3’UTR] jamIs3[Pmyo-2:DsRed::unc-54 3’UTR] jamls4[Pmyo-3::rde-4(+)::rde-4 3’UTR & pHC183)] II* (Outcrossed to AMJ8)

AMJ162 *rde-4(ne301); jamEx24[Pnas-9::rde-4(+)::rde-4 3’UTR & Pnas-9::gfp::unc-54 3’UTR]*

AMJ188 *rde-4(ne301); jamEx3[Prgef-1::rde-4(+)::rde-4 3’UTR & Prgef-1::gfp::unc-54 3’UTR]*

AMJ190 *rde-4(ne301) III; oxSi221[Peft-3p::gfp + cb-unc-119(+)] II*

AMJ210 *jamEx24[Pnas-9::rde-4(+)::rde-4 3’UTR & Pnas-9::gfp::unc-54 3’UTR]*

AMJ212 *rde-4(ne301) III; jamEx47[Pwrt-2::rde-4(+)::rde-4 3’UTR & Pwrt-2::gfp::unc-54 3’UTR]*

AMJ217 *rde-4(ne301) III; jamEx52[Punc-54::rde-4(+)::rde-4 3’UTR & Punc-54::gfp::unc-54 3’UTR]*

AMJ220 *rde-4(ne301) III; jamEx55[Psid-2::rde-4(+)::rde-4 3’UTR & Psid-2::gfp::unc-54 3’UTR]*

AMJ229 *rde-4(ne301) III; oxSi221 II; jamEx4*

AMJ237 *rde-4(ne301) III; jamEx65[Pmyo-3::rde-4(+)::rde-4 3’UTR & Pmyo-3::gfp::unc-54 3’UTR]*

AMJ238 *rde-4(ne301) III; jamEx66[Pmyo-3::rde-4(+)::rde-4 3’UTR & Pmyo-3::gfp::unc-54 3’UTR]*

AMJ239 *rde-4(ne301) III; jamEx67[Pmyo-3::rde-4(+)::rde-4 3’UTR & Pmyo-3::gfp::unc-54 3’UTR]*

AMJ268 *rde-4(ne301) III; sid-1(qt9) V; jamEx3*

AMJ269 *rde-4(ne301) III; sid-1(qt9) V*

AMJ290 *rde-4(ne301) III; jamEx89[Punc-119c::rde-4(+)::rde-4 3’UTR & Punc-119c::gfp::unc-54 3’UTR)]*

AMJ303 *jamEx77[Pnas-9::gfp::unc-54 3’UTR]*

AMJ311 *rde-4(ne301) III; sid-1(qt9) V; jamEx12*

AMJ314 *oxSi221 II; unc-119(ed3) III (?); rde-1(ne219) V*

AMJ331 *oxSi221 II; unc-119(ed3) III (?); rde-1(ne219) V; jamEx1*

AMJ343 *rde-4(ne301) III; sid-1(qt9) V; jamEx3 jamEx12*

AMJ383 *eri-1(mg366) IV; jamEx77*

AMJ385 *nrde-3(tm1116) X; jamEx77*

AMJ488 *ergo-1(tm1860) V; jamEx77*

AMJ565 *jamSi6 II; rde-4(ne301) III; unc-119(ed3) III (?)*

AMJ749 *bli-1(jam14) II*

AMJ783 *jamEx194[Pmyo-3::DsRed::unc-54 3’UTR]* – line 1

AMJ784 *jamEx195[Pmyo-3::DsRed::unc-54 3’UTR]* – line 2

AMJ785 *jamEx196[Pmyo-3::DsRed::unc-54 3’UTR]* – line 3

AMJ788 *rde-1(ne219) V; jamEx199[Psid-2::rde-1(+)::rde-1 3’UTR & Psid-2::gfp::unc-54 3’UTR]*

AMJ793 *jamEx203[Prgef-1::bli-1-dsRNA & Prgef-1::gfp::unc-54 3’UTR]*

AMJ804 *rde-4(ne301) III; K08F4.2(K08F4.2::gfp) IV*

AMJ805 *oxSi221 II; unc-119(ed3) III(?); jamEx196*

AMJ821 *nrde-3(tm1116) X; jamEx203*

AMJ822 *sid-1(qt9) V; jamEx203*

AMJ824 *rde-4(ne301) III; K08F4.2(K08F4.2::gfp) IV; jamEx4*

AMJ912 *jamSi28[Pmyo-3::rde-4(+)::rde-4 3’UTR] II; rde-4(ne301) III*

EG4322 *ttTi5605 II; unc-119(ed3) III*

EG6070 *oxSi221 II; unc-119(ed3)*

GR1373 *eri-1(mg366) IV*

HC196 *sid-1(qt9) V*

HC780 *rrf-1(ok589) I*

JH3197 *K08F4.2(K08F4.2::gfp) IV*

WM27 *rde-1(ne219) V*

WM49 *rde-4(ne301) III*

WM156 *nrde-3(tm1116) X*

WM158 *ergo-1(tm1860) V*

YY160 *nrde-1(gg88) III*

YY186 *nrde-2(gg91) II*

YY453 *nrde-4(gg129) IV*

### Transgenesis

<u>To express *rde-4(+)* in the body-wall muscle under the *myo-3* promoter:</u>

The wild-type *rde-4* gene was expressed under the control of the *myo-3* promoter from extrachromosomal arrays (AMJ66 (23), AMJ237, AMJ238, and AMJ239) or from an integrated array (AMJ151)).

Expression from extrachromosomal arrays: To make *Pmyo-3::rde-4(+)::rde-4 3’UTR*, the *myo-3* promoter (*Pmyo-3*) was amplified with primers P28 and P31, and *rde-4(+)* was amplified with primers P30 and P4. The two PCR products were used as templates to generate the *Pmyo-3::rde-4(+)* fusion product with primers P29 and P32. To make *Pmyo-3::gfp::unc-54 3’UTR*, gDNA from a strain with a transgene that expresses *Pmyo-3::gfp::unc-54 3’UTR* (HC150 (*ccIs4251 [pSAK2 (Pmyo-3::nlsGFP-LacZ) & pSAK4 (Pmyo-3::mitoGFP), & dpy-20 subclone)] I; qtIs3 [pBMW14(Pmyo-2::GFP–unc-22–PFG)] III; mIs11 [Pmyo-2::gfp, gut::gfp, pes-10::gfp] IV sid(qt25)*) was used as template to directly amplify the fusion product with the primers P33 and P34 using Long-Template Expand Polymerase (Roche). WM49 animals were microinjected with a 1:1 mixture (10 ng/μl) of *Pmyo-3::rde-4(+)::rde-4 3’UTR* and *Pmyo-3::gfp::unc-54 3’UTR* in 10 mM Tris (pH 8.5) to generate three independent transgenic lines (AMJ237, AMJ238, and AMJ239).

Expression from an integrated array: A strain with two spontaneous integration events that generated *jamIs3* and *jamIs4* was designated as AMJ151 (*rde-4(ne301) III; mIs11 jamIs3 jamIs4 IV*). Microinjection of pHC448 at 38 ng/μl in 10 mM Tris (pH 8.5) into *rde-4(ne301) III; mIs11* generated *jamIs3*. Subsequent microinjection of a mix of *Pmyo-3::rde-4* and pHC183 (as described earlier in (23)) generated *jamIs4*. The resultant strain was then outcrossed by mating with AMJ8 (*juls73*) to generate *juls73*/*rde-4(ne301)* heterozygotes and picking their self progeny that lack *juls73*.

<u>To express *rde-4(+)* in the body-wall muscle under the *unc-54* promoter:</u>

To make *Punc-54::rde-4(+)::rde-4 3’UTR*, the *unc-54* promoter (*Punc-54*) was amplified with primers P22 and P24, and *rde-4(+)* and *rde-4 3’UTR* was amplified with primers P23 and P4. The two PCR products were used as templates to generate the *Punc-54::rde-4(+)::rde-4 3’UTR* fusion product with primers P25 and P5. To make *Punc-54::gfp::unc-54 3’UTR, Punc-54* was amplified using primers P22 and P27 and *gfp::unc-54 3’UTR* was amplified from pPD95.75 using primers P26 and P8. The two PCR products were used as templates and *Punc-54::gfp::unc-54 3’UTR* fusion product was generated using the primers P25 and P13. WM49 animals were microinjected with a 1:1 mixture (10 ng/μl) of *Punc-54::rde-4(+)::rde-4 3’UTR* and *Punc-54::gfp::unc-54 3’UTR* in 10 mM Tris (pH 8.5) to generate transgenic lines. A representative transgenic line was designated as AMJ217.

<u>To express *rde-4(+)* in the hypodermis under the *nas-9* promoter:</u>

To make *Pnas-9::rde-4(+)::rde-4 3’UTR*, the *nas-9* promoter (*Pnas-9*) was amplified using the primers P1 and P3, and *rde-4(+)* and *rde-4 3’UTR* was amplified using the primers P2 and P4. The two PCR products were used as templates to generate the *Pnas-9::rde-4(+)::rde-4 3’UTR* fusion product with primers P40 and P5. To make *Pnas-9::gfp::unc-54 3’UTR, Pnas-9* was amplified with primers P1 and P7, and *gfp::unc-54 3’UTR* was amplified from pPD95.75 using the primers P6 and P8. WM49 animals were microinjected with a 2:1:1 mixture of *Pnas-9::rde-4(+)::rde-4 3’UTR* (10 ng/μl)*, Pnas-9* (with *gfp* overlap) (5 ng/μl) and *gfp* (with *Pnas-9* overlap) (5 ng/μl) in 10 mM Tris (pH 8.5) to generate transgenic lines. A representative transgenic line was designated as AMJ162.

To make strain AMJ210, AMJ162 was crossed with AMJ8 males. F2 cross progeny that were homozygous for *juls73* (which is linked to *rde-4(+)*) and contained the *jamEx24[Pnas-9::rde-4(+)::rde-4 3’UTR & Pnas-9::gfp::unc-54 3’UTR]* transgene were passaged for one generation to ensure homozygosity of *juls73* and then crossed with N2 males. A representative F2 progeny of this cross that lacked *juls73* (i.e. was homozygous for *rde-4(+)*) but contained the *jamEx24[Pnas-9::rde-4(+)::rde-4 3’UTR & Pnas-9::gfp::unc-54 3’UTR]* transgene was designated as AMJ210.

<u>To express *rde-4(+)* in the hypodermis under the *wrt-2* promoter:</u>

To make *Pwrt-2::rde-4(+)::rde-4 3’UTR*, the *wrt-2* promoter (*Pwrt-2*) was amplified using the primers P9 and P11, and *rde-4(+)::rde-4 3’UTR* was amplified using the primers P10 and P4. The two PCR products were used as templates to generate the *Pwrt-2::rde-4(+)::rde-4 3’UTR* fusion product with primers P12 and P5. To make *Pwrt-2::gfp::unc-54 3’UTR, Pwrt-2* was amplified using primers P9 and P15, and *gfp::unc-54 3’UTR* was amplified from pPD95.75 using primers P14 and P8. The two PCR products were used as templates to generate *Pwrt-2*::*gfp* fusion product with primers P12 and P13. WM49 animals were microinjected with a 1:1 mixture (10 ng/μl) of *Pwrt-2::rde-4(+)::rde-4 3’UTR* and *Pwrt-2::gfp::unc-54 3’UTR* in 10 mM Tris (pH 8.5) to generate transgenic lines. A representative transgenic line was designated as AMJ212. This strain grew slowly for the first ~4 generations, but became comparable to other strains in later generations.

<u>To express *rde-4(+)* in the intestine under the *sid-2* promoter:</u>

To make *Psid-2::rde-4(+)::rde-4 3’UTR*, the *sid-2* promoter (*Psid-2*) was amplified using the primers P16 and P18, and *rde-4(+)* along with *rde-4* 3’UTR was amplified using the primers P17 and P4. The two PCR products were used as templates and the *Psid-2::rde-4(+)::rde-4 3’UTR* fusion product was generated using the primers P19 and P5. To make *Psid-2::gfp::unc-54 3’UTR, Psid-2* was amplified using the primers P16 and P21, and *gfp::unc-54 3’UTR* was amplified from pPD95.75 using the primers P20 and P8. The two PCR products were used as templates and the *Psid-2::gfp::unc-54 3’UTR* fusion product was generated using the primers P19 and P13. WM49 animals were microinjected with a 1:1 mixture (10 ng/μl) of *Psid-2::rde-4(+)::rde-4 3’UTR* and *Psid-2::gfp::unc-54 3’UTR* in 10 mM Tris (pH 8.5) to generate transgenic lines. A representative transgenic line was designated as AMJ220.

<u>To express *rde-4(+)* in the neurons under the *rgef-1* promoter:</u>

*rde-4(ne301); qtEx136; jamEx3[Prgef-1::rde-4(+)::rde-4 3’UTR & Prgef-1::gfp::unc-54 3’UTR]* animals (described in (23)) were crossed with WM49 and *rde-4(ne301); jamEx3[Prgef-1::rde-4(+)::rde-4 3’UTR & Prgef-1::gfp::unc-54 3’UTR]* progeny were isolated and designated as AMJ188.

<u>To express *rde-4(+)* in the neurons under the *unc-119* promoter:</u>

To make *Punc-119::rde-4(+)::rde-4 3’UTR*, the *unc-119* promoter (*Punc119*) was amplified using the primers P39 and P35, and *rde-4(+)::rde-4 3’UTR* was amplified using the primers P38 and P4. The two PCR products were used as templates to generate the *Punc-119::rde-4(+)::rde-4 3’UTR* fusion product with primers P41 and P5. To make *Punc-119::gfp::unc-54 3’UTR, Punc119* was amplified using primers P39 and P36, and *gfp::unc-54 3’UTR* was amplified from pBH34.21 using the primers P37 and P42. The two PCR products were used as templates and the *Punc-119::gfp::unc-54 3’UTR* fusion product was generated using the primers P41 and P43. WM49 animals were microinjected with a 1:1 mixture (10 ng/μl) of *Punc-119::rde-4(+)::rde-4 3’UTR* and *Punc-119::gfp::unc-54 3’UTR* in 10 mM Tris (pH 8.5) to generate transgenic lines. A representative transgenic line was designated as AMJ290.

<u>To express *rde-1(+)* in the body-wall muscles under the *myo-3* promoter:</u>

As described in (23).

<u>To express *rde-1(+)* in the intestine under the *sid-2* promoter:</u>

To make *Psid-2::rde-1(+)::rde-1 3’UTR*, the *sid-2* promoter (*Psid-2*) was amplified using the primers P16 and P65, and *rde-1(+)::rde-1 3’UTR* was amplified using the primers P66 and P45. The two PCR products were used as templates to generate the *Psid-2::rde-1(+)::rde-1 3’UTR* fusion product with primers P19 and P46. Coinjection marker *Psid-2::gfp::unc-54 3’UTR*, was made as described for AMJ220. WM27 animals were microinjected with a 1:1 mixture (10 ng/μl) of *Psid-2::rde-1(+)::rde-1 3’UTR* and *Psid-2::gfp::unc-54 3’UTR* in 10 mM Tris (pH 8.5) to generate transgenic lines. A representative transgenic line was designated as AMJ788.

<u>To express *rde-1(+)* in the hypodermis under the *wrt-2* promoter:</u>

To make *Pwrt-2::rde-1(+)::rde-1 3’UTR*, the *wrt-2* promoter (*Pwrt-2*) was amplified using the primers P9 and P43, and *rde-1(+)::rde-1 3’UTR* was amplified using the primers P44 and P46. The two PCR products were used as templates to generate the *Pwrt-2::rde-1(+)::rde-1 3’UTR* fusion product with primers P12 and P46. *Pwrt-2::gfp::unc-54 3’UTR* was made as described for AMJ212. WM27 animals were microinjected with a 1:1 mixture (10 ng/μl) of *Pwrt-2::rde-1(+)::rde-1 3’UTR* and *Pwrt-2::gfp::unc-54 3’UTR* in 10 mM Tris (pH 8.5) to generate transgenic lines. Three representative transgenic lines were designated as AMJ631, AMJ632, and AMJ633.

<u>To express *rde-1(+)* in neurons under the *rgef-1* promoter:</u>

Made as described in (23).

<u>To express *gfp* in the hypodermis under the *nas-9* promoter:</u>

To make *Pnas-9::gfp::unc-54 3’UTR*, the *nas-9* promoter (*Pnas-9*) was amplified with primers P1 and P6, and *gfp::unc-54 3’UTR* was amplified from pPD95.75 using primers P47 and P8. The two PCR products were used as templates and *Pnas-9::gfp::unc-54 3’UTR* fusion product was generated using the primers P40 and P13. N2 animals were microinjected with a *Pnas-9::gfp::unc-54 3’UTR* in 10 mM Tris (pH 8.5) to generate transgenic lines. A representative transgenic line was designated as AMJ303.

<u>To express *DsRed* in the body-wall muscle under the *myo-3* promoter:</u>

N2 animals were microinjected with pHC183 (*Pmyo-3::DsRed::unc-54 3’UTR*, made as described in (23)) in 10 mM Tris (pH 8.5) to generate 3 transgenic lines designated as AMJ783, AMJ784 and AMJ785.

<u>To express *sid-1(+)* in the body-wall muscles under the *myo-3* promoter:</u>

As described in (6).

<u>To express *rde-4(+)* in the hypodermis under the *nas-9* promoter from a single-copy transgene:</u>

EG4322 animals were microinjected with a mixture of pJM6 (22.5ng/μl) and the coinjection markers pCFJ601 (50ng/μl), pMA122 (10 ng/μl), pGH8 (10 ng/μl), pCFJ90 (2.5 ng/μl), and pCFJ104 (5 ng/μl) (plasmids described in (27)) to generate a transgenic line as described earlier (27). This isolated line was crossed into AMJ8 males and the resulting *rde-4(+)/juls73* male progeny were crossed to WM49, and homozygozed for the single-copy insertion and *rde-4(-)* to generate AMJ565. The integration of *Pnas-9::rde-4(+)::rde-4 3’UTR* in AMJ565 was verified by genotyping for the presence of *Pnas-9::rde-4(+)* using primers P53 and P54.

<u>To express *rde-4(+)* in the body-wall muscle under the *myo-3* promoter from a single-copy transgene:</u>

EG4322 animals were microinjected with a mixture of pPR1 (22.5ng/μl) and the coinjection markers pCFJ601 (50ng/μl), pMA122 (10 ng/μl), pGH8 (10 ng/μl), pCFJ90 (2.5 ng/μl), and pCFJ104 (5 ng/μl) (plasmids described in (27)) to generate a transgenic line as described earlier (27). This isolated line was crossed into AMJ8 males and the resulting *rde-4(+)/juls73* male progeny were crossed to WM49, and homozygozed for the single-copy insertion and *rde-4(-)* to generate AMJ912. The integration of *Pmyo-3::rde-4(+)::rde-4 3’UTR* in AMJ912 was verified by genotyping for the presence of an insertion at the Mos site using primers P82-P84.

<u>To express *bli-1-dsRNA* in the neurons under the *rgef-1* promoter:</u>

To make *Prgef-1::bli-1-dsRNA* sense strand, the *rgef-1* promoter (*Prgef-1*) was amplified with primers P67 and P68 and a 1kb region in exon 3 of *bli-1* was amplified using primers P69 and P70. The two PCR products were used as templates and *Prgef-1::bli-1-dsRNA* sense fusion product was generated using the primers P71 and P72. To make *Prgef-1::bli-1-dsRNA* antisense strand, the *rgef-1* promoter (*Prgef-1*) was amplified with primers P67 and P73 and *bli-1* was amplified using primers P74 and P75. The two PCR products were used as templates and *Prgef-1::bli-1-dsRNA* antisense fusion product was generated using the primers P71 and P76. N2 animals were microinjected with a 1:1:1 ratio of sense *Prgef-1::bli-1-dsRNA*, antisense *Prgef-1::bli-1-dsRNA* and *Prgef-1::gfp::unc-54 3’UTR* (as described in (23)) in 10 mM Tris (pH 8.5) to generate transgenic lines. A representative transgenic line was designated as AMJ793.

### SUPPLEMENTAL FIGURES AND FIGURE LEGENDS

**Figure S1.**
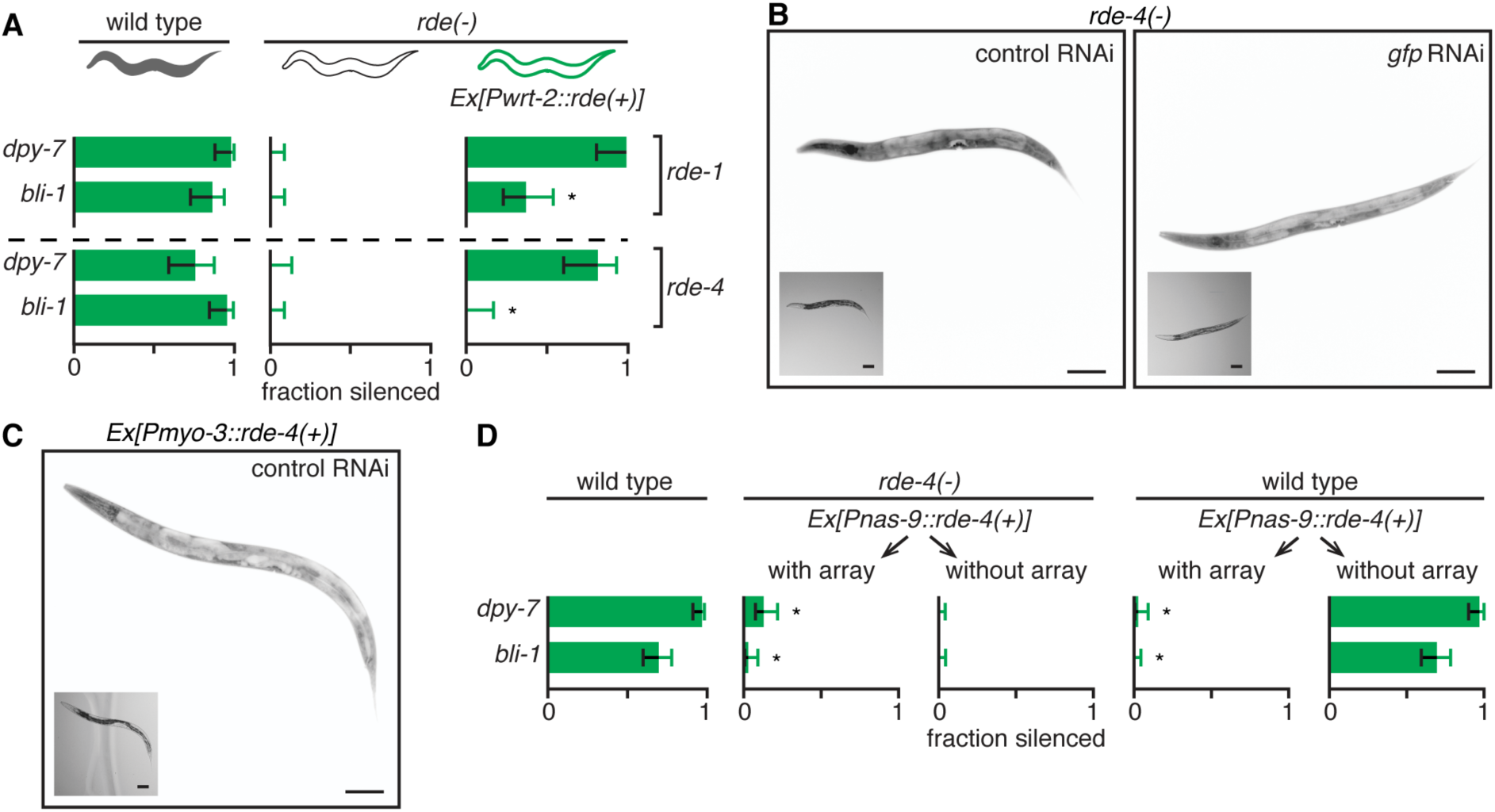
Silencing by feeding RNAi of some genes is reduced in tissues expressing RDE-4 or RDE-1 from a repetitive transgene. (**A**) Silencing by feeding RNAi of some endogenous genes is reduced in tissues expressing *rde-4(+)* or *rde-1(+)* from a repetitive transgene. Wild-type animals, mutant animals (*rde-1(-)*, top or *rde-4(-)*, bottom) or mutant animals with tissue specific rescue in the hypodermis (*Ex[Pwrt-2::rde(+)]*) were fed dsRNA against *dpy-7* or *bli-1* and the fractions of animals that showed silencing (fraction silenced) were determined. (**B** and **C**) Representative images of animals that express *gfp* in all somatic cells (*Peft-3::gfp*) in a *rde-4(-)* background (B) or with *rde-4* rescued in the muscles (*Pmyo-3::rde-4(+)*) (C) that were fed either control dsRNA (control RNAi) or dsRNA against *gfp* (*gfp* RNAi) are shown. Insets are brightfield images, scale bar = 50 μm. Also see Figure 1B. (**D**) Silencing of *bli-1* and *dpy-7* is inhibited by the expression of RDE-4 from a repetitive transgene in the hypodermis. Wild-type animals and *rde-4(-)* animals that express *rde-4(+)* along with *gfp* in the hypodermis (*Ex[Pnas-9::rde-4(+)]*) from extrachromosomal repetitive DNA (array) were fed dsRNA against hypodermal genes (*dpy-7* or *bli-1*, green). The fractions of animals either with or without the arrays that showed silencing (fraction silenced) were determined. Also see Figure 1C. Error bars indicate 95% confidence intervals (CI), n>22 animals and asterisks indicates p<0.01.

**Figure S2.**
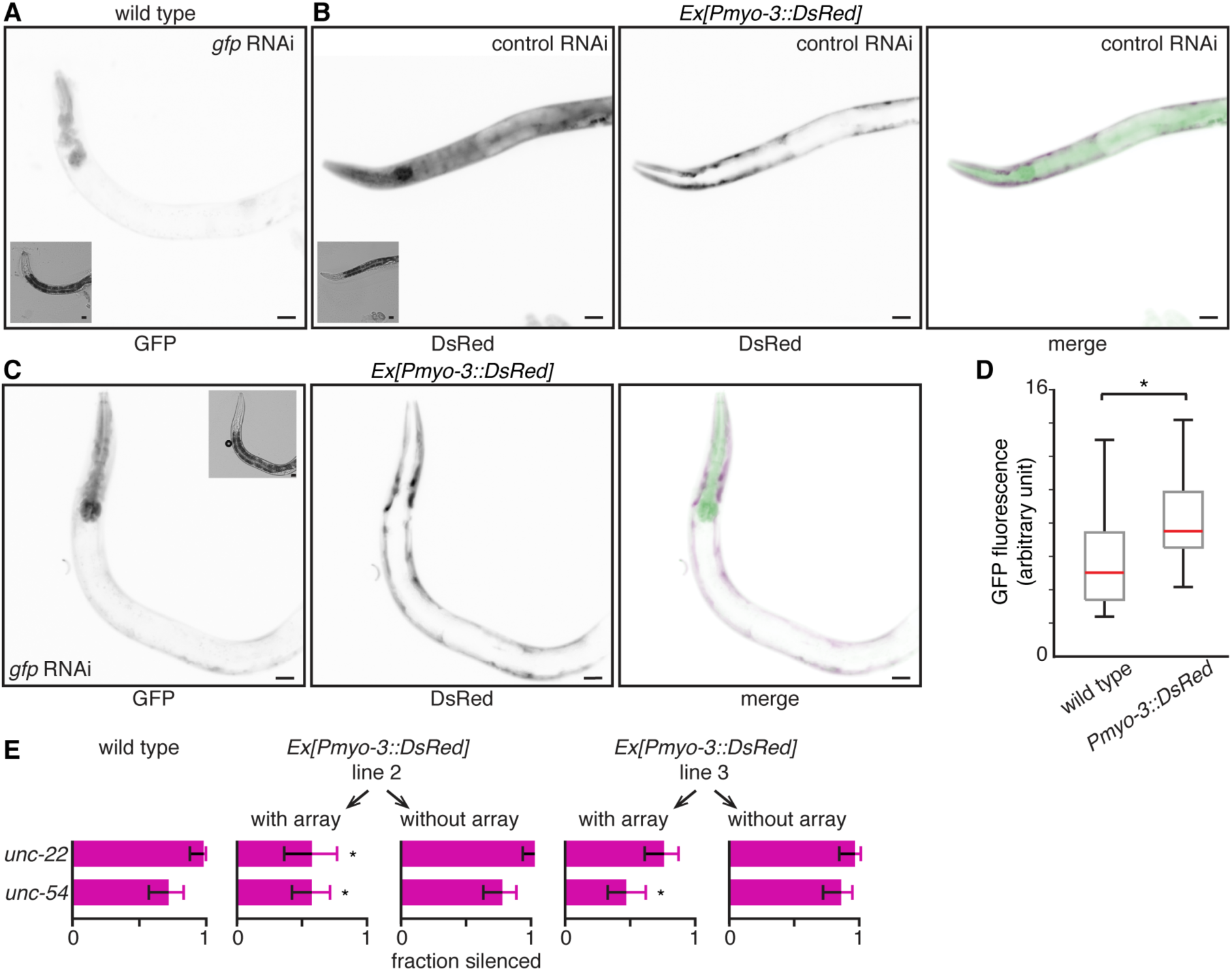
Silencing by feeding RNAi of some genes is reduced in tissues expressing a repetitive transgene of unrelated sequence. (**A**-**C**) Representative images of animals with *gfp* expression in all somatic cells (GFP) that were fed *gfp* RNAi (A) or animals that in addition express *DsRed* in the body-wall muscle (*Pmyo-3::DsRed*) that were fed control RNAi (B) or *gfp* RNAi (C) are shown. Merged images in (C) show overlap of *gfp* and *DsRed* expression (DsRed = magenta and GFP = green). Insets are brightfield images and scale bar = 50 μm. (**D**) Quantification of *gfp* silencing in animals with or without *DsRed* expression in body-wall muscles. In animals expressing *DsRed* in body-wall muscles (*Ex[Pmyo-3::DsRed]*), *gfp* fluorescence was brighter in body-wall muscles than in other tissues. Red lines indicate median GFP fluorescence. (**E**) Silencing of *unc-22* and *unc-54* can be inhibited by the expression of a repetitive transgene of unrelated sequence in body-wall muscles. Two additional lines (also see Figure 1D) of wild-type animals expressing *DsRed* in body-wall muscles (*Ex[Pmyo-3::DsRed]*) from extrachromosomal repetitive transgenes (array) were fed dsRNA against body-wall muscle genes (*unc-22* or *unc-54*, magenta). The fractions of animals either with or without the arrays that showed silencing (fraction silenced) were determined (n=50). Error bars indicate 95% CI and asterisks indicate p<0.01.

**Figure S3.**
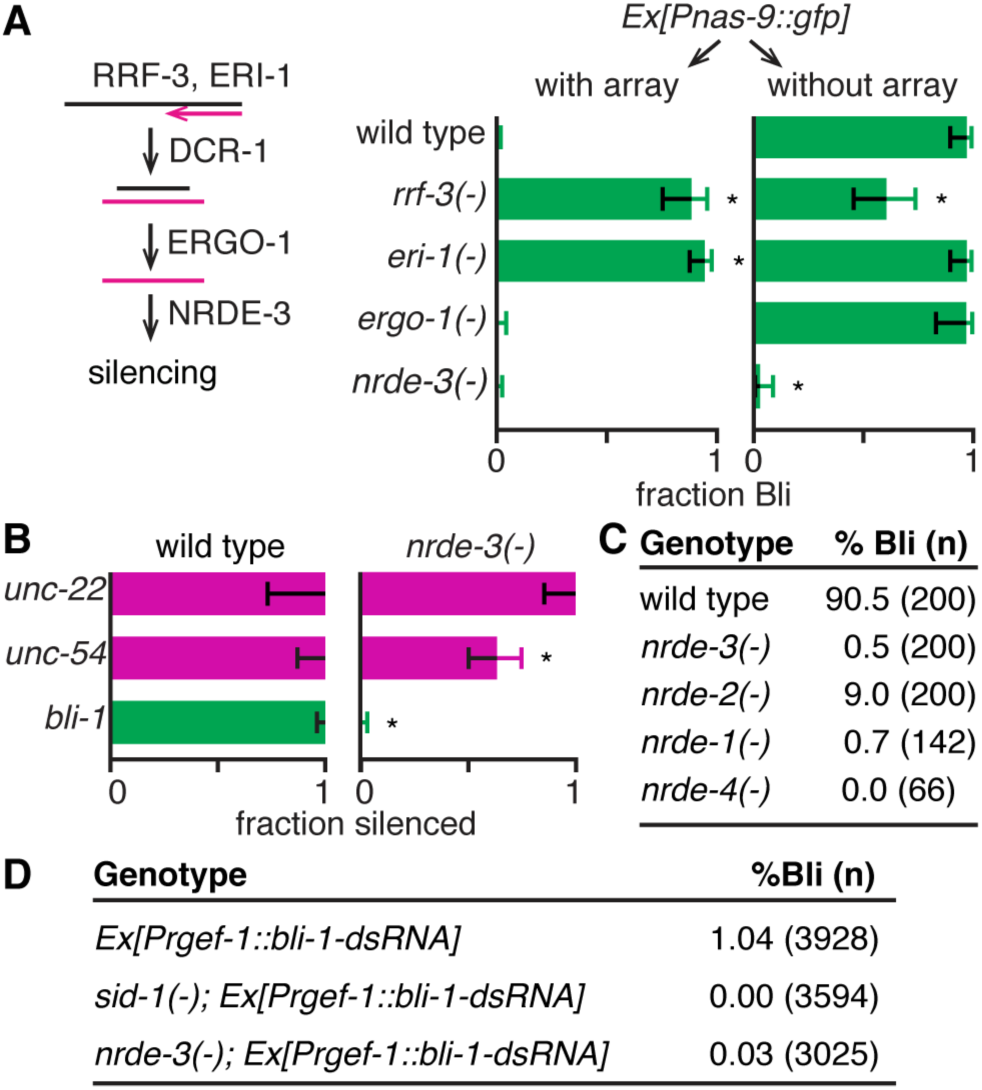
Genes that show robust inhibition of silencing upon expression of an unrelated transgene can recruit the nuclear RNAi pathway for silencing. (**A**) Inhibition of *bli-1* silencing by expression of any repetitive DNA can be relieved by the loss of some components of the endogenous RNAi pathway. (Left) Schematic of endogenous RNAi. Aberrant RNA recruits the RNA-dependent RNA polymerase RRF-3, the exonuclease ERI-1, the endonuclease DCR-1, the primary Argonaute ERGO-1, and the secondary Argonautes (e.g. NRDE-3) to cause silencing. (Right) Loss of *eri-1* or *rrf-3* but not of *ergo-1* or *nrde-3* relieves inhibition of *bli-1* silencing caused by hypodermal expression of repetitive DNA. Extent of silencing (fraction Bli) in response to *bli-1* feeding RNAi of animals with or without an extrachromosomal array that expresses *gfp* in the hypodermis (*Ex[Pnas-9::gfp]*) in a wild-type, *rrf-3(-), eri-1(-), ergo-1(-)* or *nrde-3(-)* background were determined (n>28 animals). (**B**) Genes that show robust inhibition of silencing upon expression of a repetitive transgene appear to also require the nuclear Argonaute NRDE-3 for complete silencing. Wild-type animals and *nrde-3(-)* animals were fed dsRNA against *unc-22, unc-54*, or *bli-1* and the fractions of animals that showed silencing (fraction silenced) were determined. (**C**) Silencing of *bli-1*, by ingested dsRNA requires multiple components of the nuclear RNAi pathway. Wild-type, *nrde-3(-), nrde-2(-), nrde-1(-)*, or *nrde-4(-)* animals were fed dsRNA against *bli-1* and the percentage of gravid adult animals that showed silencing (% Bli) was determined (n = number of gravid adults scored for silencing). (**D**) Silencing of *bli-1* by neuronal dsRNA requires the nuclear Argonaute *nrde-3*. Silencing of *bli-1* by dsRNA against *bli-1* expressed under a neuronal promoter (*Ex[Prgef-1::bli-1-dsRNA]*) in a wild-type, *sid-1(-)*, or *nrde-3(-)* background were measured as in (C). Error bars indicate 95% CI and asterisks indicate p<0.01.

**Figure S4.**
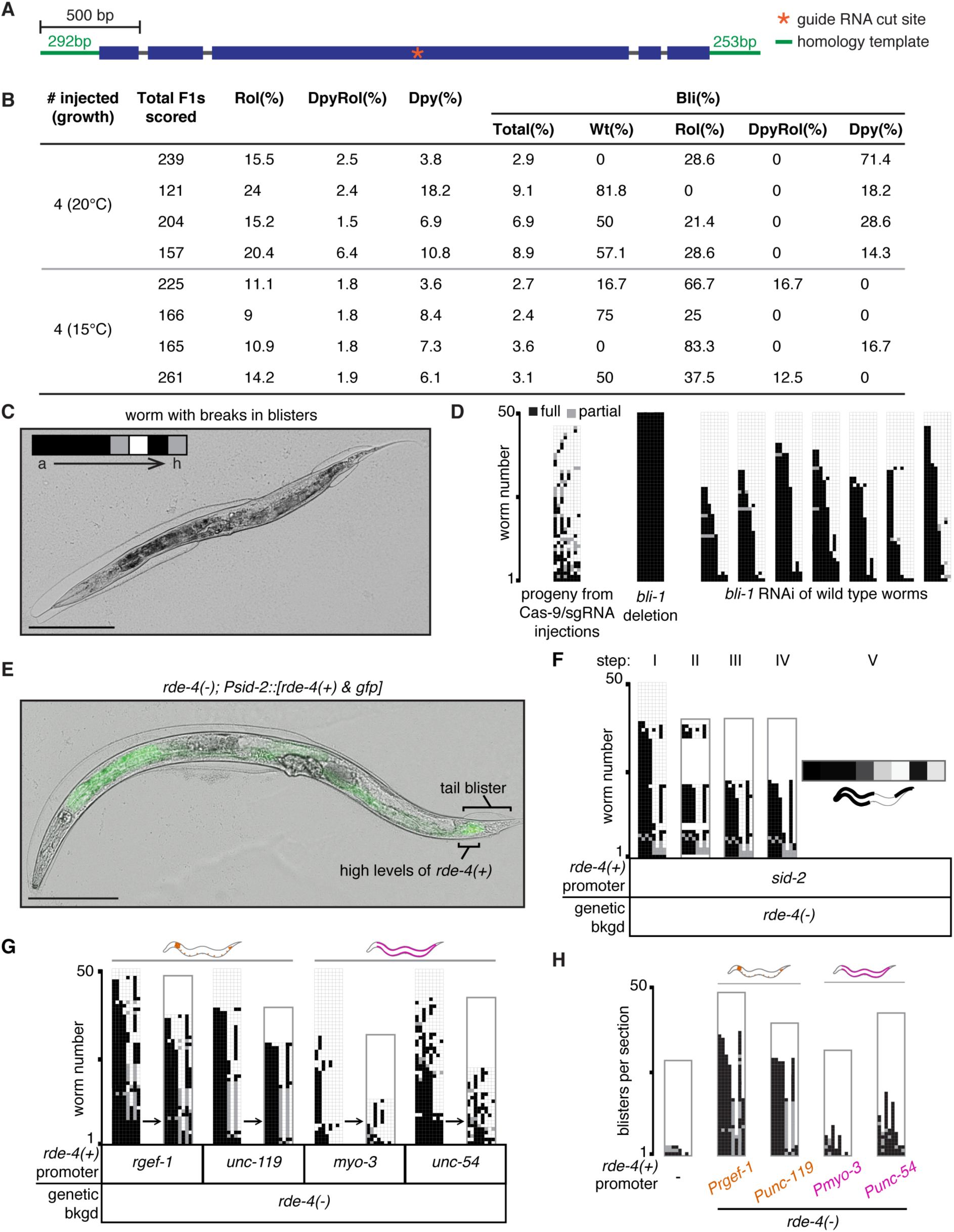
Silencing of *bli-1* upon feeding RNAi in wild-type animals and animals with tissue-specific rescue of *rde-4* in non*-*hypodermal cells results in unique patterns of blisters that differ from those in *bli-1(-)* animals. (**A**) Design of Cas9-based genome editing to generate a *bli-1* null mutant. The *bli-1* gene (exons, blue boxes; introns, grey lines) was targeted by a single-guide RNA (sgRNA) that cuts within the gene (orange *) and was repaired with a double-stranded DNA template (green flanking the gene). (**B**) Results from Cas9/sgRNA injection into wild-type worms to generate a *bli-1* null mutant. Wild-type animals were simultaneously targeted for edits in *dpy-10* and *bli-1* (co-conversion strategy (28)), resulting in progeny that displayed Rol, DpyRol and Dpy defects (indicative of *dpy-10* editing) with or without blisters as indicated. (**C**) A representative animal that illustrates scoring of *bli-1* silencing in each section of the worm. The animal was divided into 8 sections (a through h, see Figure 3C) and each section was scored for presence of a full blister (black), partial blister (grey), or no blister (white) as indicated in the inset. Scale bar = 50 μm. (**D**) Wild-type animals display a stereotyped pattern of susceptibility to *bli-1* feeding RNAi. Progeny of wild-type animals targeted by Cas9-based genome editing, *bli-1* null mutant animals, and wild-type animals exposed to feeding RNAi were scored for blister patterns as described in Figure 3C (n>45 gravid adult animals). Unlike sections in the progeny of animals that were injected with Cas9/sgRNA or in *bli-1* null mutants, sections in wild-type animals that were subject to *bli-1* feeding RNAi showed a stereotyped frequency of blister formation (a > b > … > h). (**E**) RDE-4 expression in intestinal cells near the tail correlates with blister formation in *rde-4(-)* hypodermis near the tail upon *bli-1* RNAi. (**F** and **G**) Blister scoring method to detect variations in the pattern of blister formation upon *bli-1* feeding RNAi. (F) The pattern of blister formation in *rde-4(-)* animals with *rde-4(+)* expressed in the intestine (*Psid-2*) was examined (step I) and animals following the stereotyped order of susceptibility to *bli-1* RNAi (a > b > … > h) were removed (step II). The remaining animals, which show variant susceptibility to *bli-1* feeding RNAi, were aggregated (step III and IV) and collapsed into a heat map (step V) to assess frequency of variant blisters in each section. Grey bounding box (step II-step IV) indicates the total number of worms that showed blister formation in each strain (n = 50 gravid adult animals) (G) The pattern of *bli-1* silencing in *rde-4(-)* animals with RDE-4 expressed from neuronal promoters (*Prgef-1 & Punc-119*) or body-wall muscle promoters (*Pmyo-3 & Punc-54*) were examined and variant blisters were isolated as in (F) (n = 50 adult animals). (**H**) The pattern of blisters that result from silencing of *bli-1* in *rde-4(-)* hypodermis are characteristic of the tissue that expresses RDE-4. Patterns of blister formation in response to ingested *bli-1* RNAi were examined in wild-type animals or in *rde-4(-)* animals that express RDE-4 under neuronal promoters (*Prgef-1* or *Punc-119*, orange) or under body-wall muscle promoters (*Pmyo-3* or *Punc-54*, magenta). For each strain, sections with full (black) or partial (grey) blister formation in animals that showed a variation in the order of susceptibility (i.e. were not a > b > … > h) were plotted. Grey bounding box and n are as in (F).

**Figure S5.**
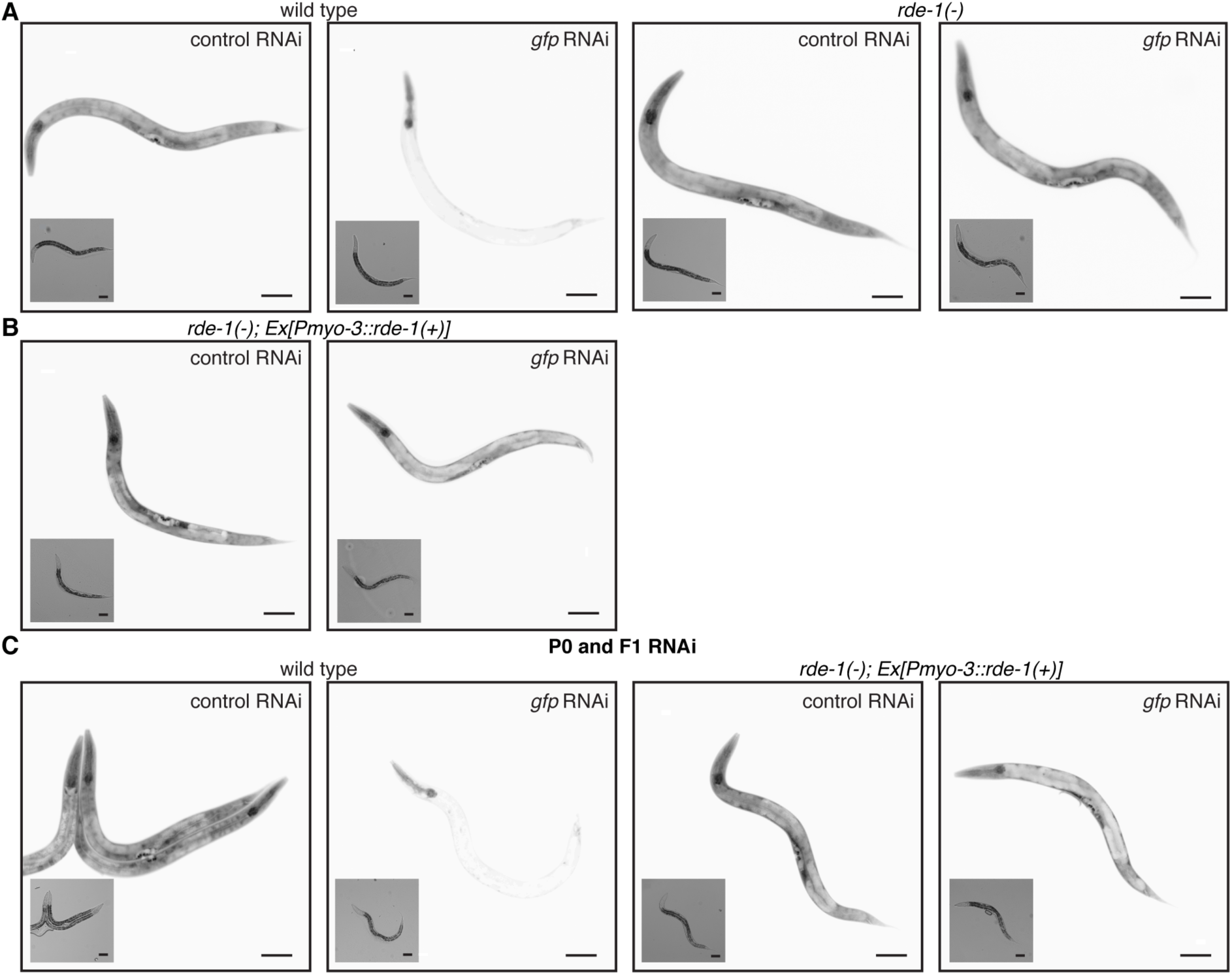
Expression of RDE-1 in one somatic tissue can enable silencing of *Peft-3::gfp* in other mutant somatic tissues. Representative images of animals with *gfp* expression in all somatic cells (*Peft-3:: gfp*) in a wild-type background (A and C), *rde-1(-)* background (A), or *rde-1(-)* background with *rde-1(+)* expressed in body-wall muscles (*Ex[Pmyo-3::rde-1(+)*) (B and C). Animals were fed control RNAi or *gfp* RNAi for one generation (A & B) or for two generations (C). Insets are brightfield images and scale bar = 50 μm.

**Figure S6.**
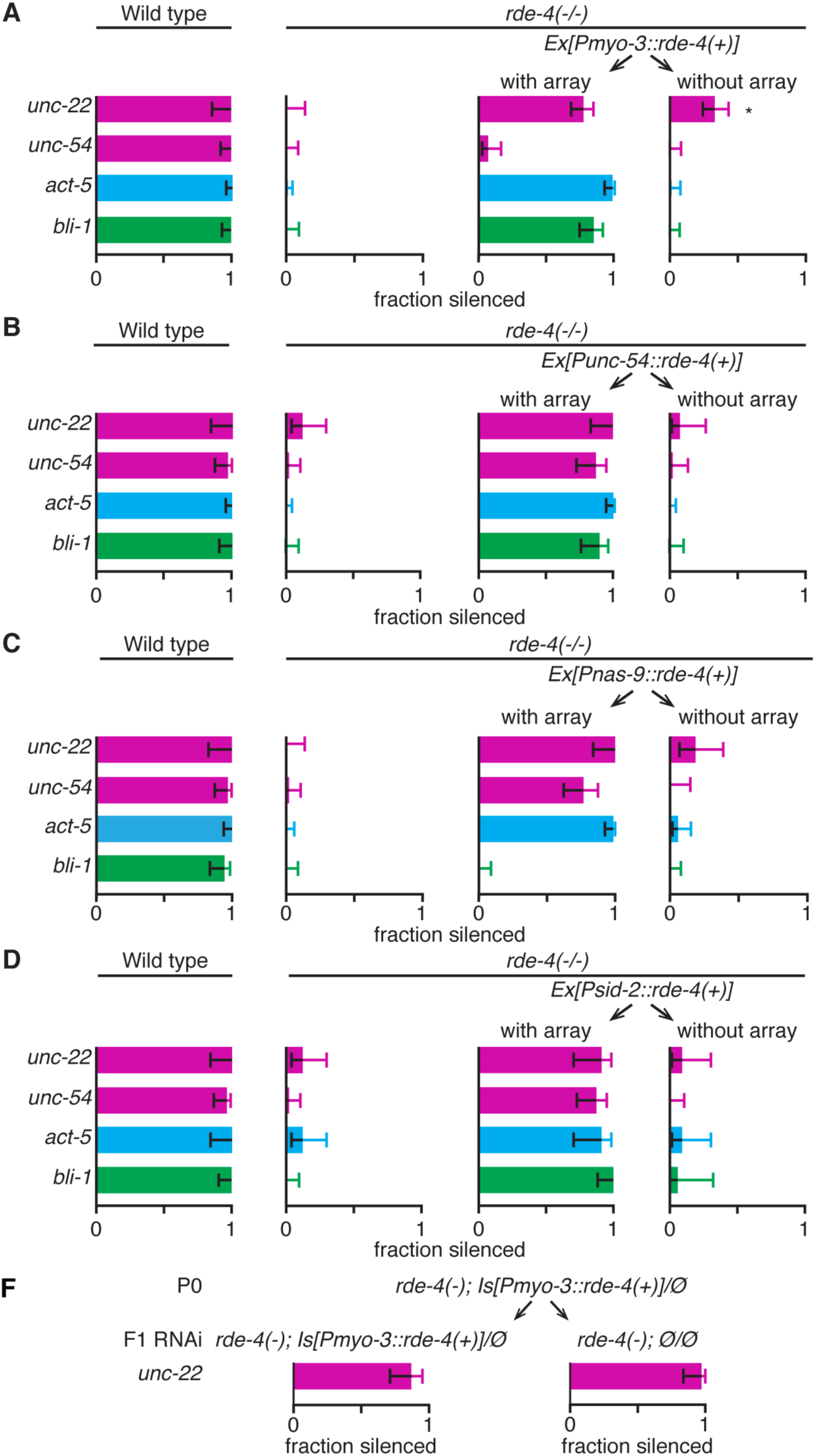
Expression of RDE-4 in the somatic tissues of parents does not typically enable feeding RNAi in *rde-4(-)* progeny. (**A**-**D**) Progeny of wild-type animals, *rde-4(-)* animals, or *rde-4(-)* animals expressing RDE-4 in the muscle *(Pmyo-3* or *Punc-54)*, hypodermis *(Pnas-9)*, or intestine *(Psid-2)* were fed dsRNA (F1-only feeding RNAi) against *unc-22, unc-54, act-5*, or *bli-1* and the fractions of animals that showed silencing (fraction silenced) were determined. **(E)** Parental expression of RDE-4 from an integrated *Pmyo-3::rde-4(+)* transgene also enables feeding RNAi of *unc-22* in *rde-4(-)* progeny. The *rde-4(+)* progeny *(Is[Pmyo-3::rde-4(+)]/Ø)* and *rde-4(-)* progeny (*Ø/Ø*) of parent animals expressing RDE-4 from an integrated repetitive array in the muscle *(Is[Pmyo-3::rde-4(+)]/Ø))* were subjected to feeding RNAi (F1 RNAi) of *unc-22* and the fractions of animals that showed silencing were determined (fraction silenced). 100% of wild-type and 0% of *rde-4(-)* control animals showed silencing of *unc-22* (n>15 animals). Error bars indicate 95% CI and asterisks indicate p<0.01.

**Figure S7.**
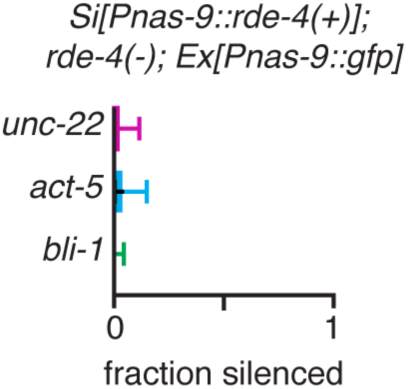
Co-expression of an unrelated gene from a repetitive transgene with RDE-4 from a single-copy transgene is insufficient to silence genes in *rde-4(-)* somatic tissues. Animals that express RDE-4 from a single-copy transgene in the hypodermis (*Si[Pnas-9::rde-4(+)]*) and additionally *gfp* from a repetitive transgene (*Ex[Pnas-9::gfp]*) in the hypodermis were fed dsRNA against *unc-22* (magenta), *act-5* (blue) or *bli-1* (green). Silencing was scored as in Figure 2A. Also see Figure 4C. Error bars indicate 95% CI and n>49 animals.

### SUPPLEMENTAL TABLES

**Table S1.**
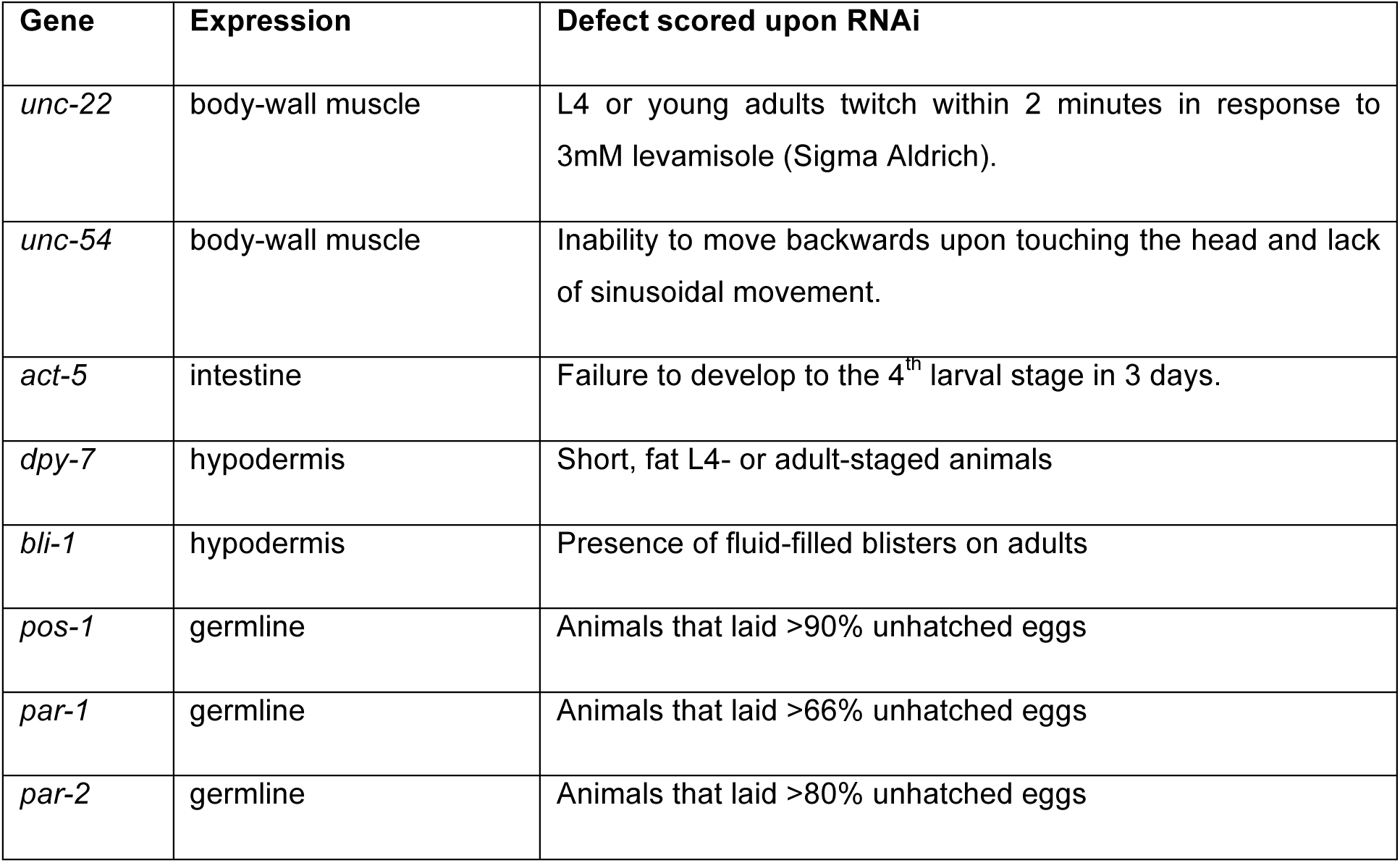
Feeding RNAi and defects scored.

| Gene | Expression | Defect scored upon RNAi |
| --- | --- | --- |
| *unc-22* | body-wall muscle | L4 or young adults twitch within 2 minutes in response to 3mM levamisole (Sigma Aldrich). |
| *unc-54* | body-wall muscle | Inability to move backwards upon touching the head and lack of sinusoidal movement. |
| *act-5* | intestine | Failure to develop to the 4<sup>th</sup> larval stage in 3 days. |
| *dpy-7* | hypodermis | Short, fat L4- or adult-staged animals |
| *bli-1* | hypodermis | Presence of fluid-filled blisters on adults |
| *pos-1* | germline | Animals that laid >90% unhatched eggs |
| *par-1* | germline | Animals that laid >66% unhatched eggs |
| *par-2* | germline | Animals that laid >80% unhatched eggs |

**Table S2.**
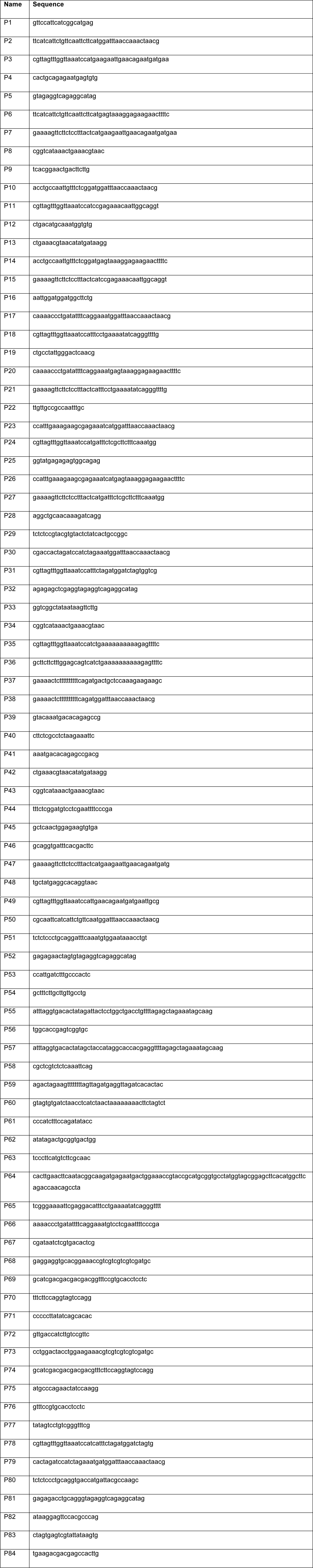
Oligonucleotides used.

